# Fast yet force-effective mode of supracellular collective cell migration on aligned fibers due to extracellular force transmission

**DOI:** 10.1101/2022.10.22.513348

**Authors:** Amrit Bagchi, Bapi Sarker, Jialiang Zhang, Marcus Foston, Amit Pathak

## Abstract

Cell collectives, like other motile entities, generate and use forces to move forward. Here, we ask whether environmental configurations alter this proportional force-speed relationship, since aligned extracellular matrix fibers are known to cause directed migration. We show that aligned fibers serve as active conduits for spatial propagation of cellular mechanotransduction through matrix exoskeleton, leading to efficient directed collective cell migration. Epithelial (MCF10A) cell clusters adhered to *soft* substrates with aligned collagen fibers (AF) migrate faster with much lesser traction forces, compared to random fibers (RF). Fiber alignment causes higher motility waves and transmission of normal stresses deeper into cell monolayer while minimizing shear stresses and increased cell-division based fluidization. By contrast, fiber randomization induces cellular jamming due to breakage in motility waves, disrupted transmission of normal stresses, and heightened shear driven flow. Using a novel motor-clutch model, we explain that such ‘force-effective’ fast migration phenotype occurs due to rapid stabilization of contractile forces at the migrating front, enabled by higher frictional forces arising from simultaneous compressive loading of parallel fiber-substrate connections. We also model *’haptotaxis*’ to show that increasing ligand connectivity (but not continuity) increases migration efficiency. According to our model, increased rate of front stabilization via higher resistance to substrate deformation is sufficient to capture ‘*durotaxis’*. Thus, our findings reveal a new paradigm wherein the rate of leading-edge stabilization determines the efficiency of supracellular collective cell migration.

## Introduction

Collective cell populations in response to various physical and chemical cues undergo directional migration that drives critical steps in development, repair, and disease [1], [2]. Cells within a collective population undergo migration in two broad phenotypes, which can operate concurrently or exclusively. First, in the presence of stable cell-cell junctions, cells behave as coupled systems driven by dynamic interplay between the active forces at the boundaries and the reciprocal coupling forces between neighboring cells [3]–[6]. Secondly, with weak or absent cell-cell junctions, cells can behave as nematic-like autonomous bodies that move in response to mutual collisions and self-correction of directionality [7]–[10]. A parameter for distinguishing these different forms of collective migration is the length scale of front-rear polarity [11]. For example, cell systems resembling active nematic would be driven by individual front-rear polarity [7]–[10]. By contrast, a collective system of coupled cells consists of two distinct populations, one of leader cells of defined front-rear polarity generating active forces and other of passive follower cells lacking polarity [12]–[14]. Here, the extent of polarity can be defined in terms of the length scale of transmission of normal stresses propagating from the leading edge into the monolayer, which is attributed to physical intercellular communication via cell-cell junctions [15]. In the event of disrupted intercellular transmission, residual stresses arise, and their shear components determine the direction of flow [16], leading to rich jamming-unjamming transitions [17], antiparallel flows [18], vortices [19] and swirls [13] in migrating epithelia. More recently, a new mode of directional migration has been demonstrated in neural crest cells of *Xenopus* [20]; here, grouped cells show extraordinary supracellular directionality due to continuity in cytoskeletal structures across coupled cells. However, the force-based mechanism behind supracellular migration is yet to be understood. More specifically, since long-range force transmission is crucial for directed collective migration, it remains an important gap in knowledge how force propagation regulates supracellular migration, which is arguably a macroscale mode of directed migration. As such, experimental and modeling frameworks connecting force generation to overall efficiency of collective cell migration remain incomplete.

Extracellular matrix fibers in tissues and organs provide biochemical and structural support to the cellular constituents. Single cells can use these fibers to sense, transmit forces and receive feedback from soft basal matrix stiffness up to 10 µm away from their immediate vicinity to modulate their migration characteristics [21]. Meanwhile, cell collectives can also self-generate aligned fibers from non-aligned structures and use them for long-range force sensing to communicate with other cells (250-1000 µm apart) [22]. As a result, aligned fibers promote long-range directed cell migration observed in cancer metastasis [23], [24], wound healing [25], [26], branching morphogenesis, and angiogenesis [27], [28]. Given the extraordinary length-scales of signal transduction and migrational directionality on aligned fibers, it is likely that cell collectives utilize supracellular modes of migration. According to previous studies, matrix stiffness and chemokine gradient regulate directional migration due to rise in migratory persistence via increased focal adhesion strength [29] and chemotaxis via increased migration-front stability [30], [31]. However, it is unknown how aligned topographies of matrix fibers, with constant stiffness or chemokine conditions, regulate directional migration. This is partly because of the inability to accurately quantify cellular migration and forces in 3D, and it remains difficult to devise reductionist soft 2D matrices with surfaces coated with aligned fibers where force measurements are viable.

Using modified polyacrylamide gels capable of attaching aligned fibers, we show that cell clusters apply lesser forces yet migrate faster using a supracellular mode of migration. In this mode, the entire monolayer behaves like a single giant polarized unit with cellular contraction occurring up to 500 µm away from the leading edge at the monolayer middle reminiscent of polarized migration of single cells. This migration phenotype is mediated by a global reduction in shear stress and an increase in fluidization. Using a novel motor-clutch model, we show that ligand engagement upon pulling of aligned fibers increases local friction, which in turn reduces traction forces and gives rise to higher cell velocity and overall migration efficiency.

## Results

### Aligned collagen fibers on soft hydrogels enhance collective cell migration speed and persistence

To probe cellular force propagation through matrix fibers, we aligned collagen-1 fibers to serve as directional cues for cell migration on soft 2D surfaces. We synthesized modified soft polyacrylamide (mod-PA) hydrogels that enable attachment of tunable collagen fibers [32], incubated in 0.1 mg/ml collagen-1, and exposed 11T magnetic field (16 hours) during collagen fibrillogenesis at 4° C (Fig. S1a) [33], resulting in aligned long collagen fibers (AF) on soft gels (2.4 kPa). Using magnetic polydimethylsiloxane (PDMS) stencils, we micro-patterned rectangular clusters (500 µm wide) of human mammary epithelial cells (MCF-10A) migrating parallel to collagen alignment (Figs. S1b-c, S1d-e). For comparison, gels with un-aligned short, random fibers (RF) were used (Fig. S1). To better understand large-scale spatiotemporal fluctuations of mechanical patterns within the expanding monolayer, we averaged velocities, tractions, and monolayer stresses across length (y-coordinate), thereby reducing dimensionality of system to a single spatial dimension (x-axis, parallel to collagen alignment) and time (Figs. 1e-h).

**Figure 1.**
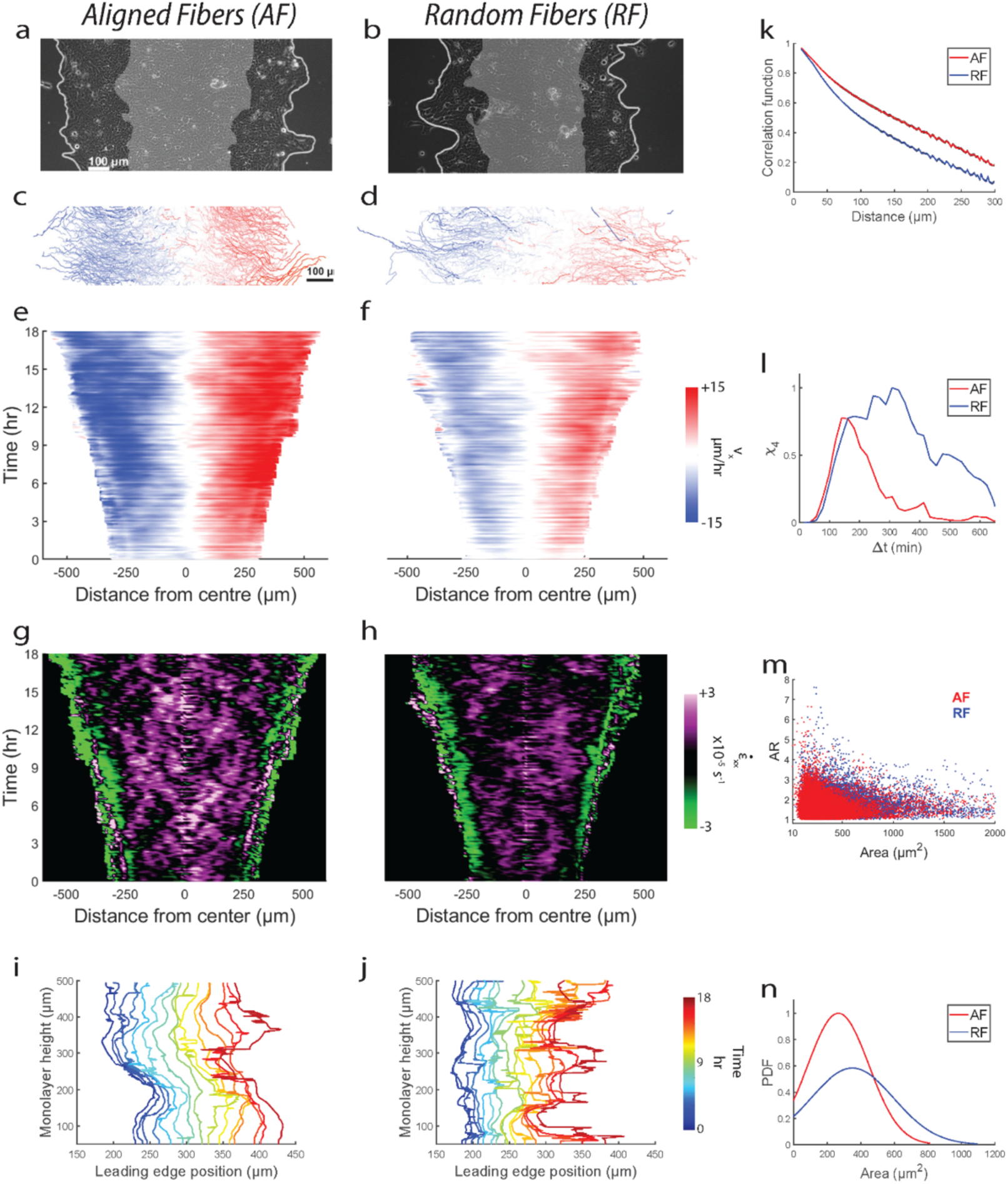
Aligned fibers cause faster and directed cell motion. Representative epithelial monolayer expanding over 16hr (gray region indicates initial position, t=0 hr) on gels with (**a**) aligned fibers (AF) and (**b**) random fibers (RF). (**c,d**) Corresponding individual cell trajectories, with color-coded mean cell speed in x-direction. Kymographs of cell velocity component in x-direction (*v_x_*) for AF (**e**) and RF (**f**), Kymographs for velocity strain rate 𝜀̇_*x*_ for AF (**g**) and RF (**h**). Plot showing temporal evolution of the leading-edge (average of left and right edges) contour for monolayer migrating on AF (**i**) and RF (**j**). (**k**) Plot comparing time-averaged (m=157) spatial autocorrelation function of 𝑣_*x*_ for AF (red) and RF (blue) for n=3. (**l**) plot comparing four-point susceptibility 𝜒_4_ versus Δ𝑡 between AF and RF. (**m**) Scatter plots of aspect ratio (AR) versus area for cells on AF (n=56117) and RF (n=39447). (**l**) Plot comparing cellular area distribution on AF and RF.

We found greater cell migration on AF over the entire 18-hour duration of migration (compared to RF; Fig. 1a,b, Fig. S2a,b). By comparing cell migration trajectories (Figs. 1c-d), we observed that cell clusters on AF migrated faster (*1.56 times) compared to RF. Additionally, migration persistence along x-axis was higher on AF (by 23%) (Fig. S1c), indicating that cells use aligned fibers as directional cues to guide their migration towards greater collective expansion. Velocity kymographs suggested cell movements for AF were more homogenous with higher velocities at the leading edge penetrating greater depths within the monolayer, compared to higher heterogeneity in RF characterized by outward faster velocities and middle regions of lesser velocities (Fig. 1e,f). Quantitatively, we measured the spatial velocity autocorrelation function 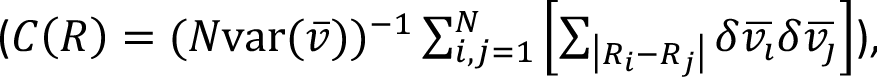, which follows a slower spatial decay in AF (correlation length higher by 26%), confirming that cellular motion is correlated across greater distances when compared to RF (Fig. 1k). We also studied the temporal evolution of the monolayer front over time for both AF and RF (Fig. 1i,j). We found greater spatial overlap of the leading front (along the width, x direction) on RF compared to AF where the front maintains its shape (along the height, y direction) across the duration of migration. Thus, the leading front on AF is more temporally stable and smoother in shape compared to RF. Consistent with recent studies connecting migration front stability to a chemotaxis-like directional migration [30], [31], we show that aligned fibers also enhance migration front stability and cause faster directed migration than random fibers.

### Collagen fiber alignment enables efficient long-range transmission of unjammed collective cell migration

To investigate temporal heterogeneity in cell movements across monolayer, we use the four-point susceptibility χ_4_, a soft matter physics tool for measuring the rate of structural rearrangements between any two points in space within a time window. We computed χ_4_ from the variance of self-overlap order parameter 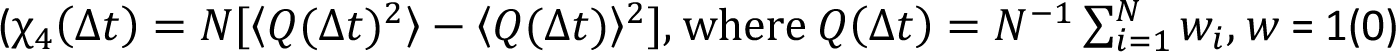 if 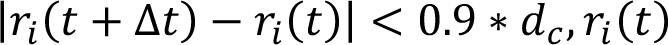 is x − position of cell at time 𝑡 and 𝑑_*c*_ is average cell diameter). While the χ_4_ peak location indicates time interval for heterogeneous cell migration dynamics, the peak height denotes spatial extent of such heterogeneities [34]. Comparing AF to RF, there is a distinct shift in χ_4_ peak to longer timescales for RF (Fig. 1j), indicating higher cellular jamming on RF. Additionally, higher peak height for RF indicates larger clusters of jammed cells. According to previous work on inert particulate matter [35], [36], differences in mutual crowding between two conditions could be responsible for distinct jamming states. Compared to AF, cells on RF had much higher area (31% higher) (Fig. 1 k,l) and slightly higher aspect ratio (2.3% higher) (Figs. 1k, S2e) along with much lesser densities (33% lower) (Fig. S2 d). Although RF cellular morphologies agree with conventional jammed state [37], reduced crowding is counter-intuitive because jamming tends to occur at higher densities in inert systems [36], [38]. Based on this experimental evidence, we decided to explore beyond the density-based jamming hypothesis to understand how fiber alignment regulates migration efficiency. It is possible that owing to smaller sizes, cells on AF find it easier to slip past each other despite higher densities. In comparison, on RF, relatively larger cell sizes coupled with uncorrelated migration in x-and y-direction can lead to higher impinging among cells, thus biasing the phenotype towards jamming [39]. For cells on AF, our analysis suggests a flocking-like migratory phenotype [40], [41], which is spatially homogenous and temporally dynamic. By contrast, on RF, migration is spatially more heterogeneous and static. We will investigate the cause and implications of cell density differences between RF and AF cases later in the manuscript, along with the underlying cause of cellular flocking/jamming on respective fiber alignment conditions.

Interestingly, for AF alone, there were multiple strain-rate X-waves [42] propagating away from leading edge, coalescing in the monolayer middle, and then propagating back to edges (Figs. 1g-h). These X-waves could not be well defined for RF. We measured the strain-rate fluctuations at a 100 µm wide region at the monolayer midline (shown in Fig. 4c-d and discussed ahead in more detail). We observed two peaks for AF inside first 10 hours of migration compared to a single peak on RF which occurred at the 10^th^ hour time point. Given the velocity and cell density differences across AF and RF, X-waves travel faster, more frequently and across greater cell-cell junctions on AF, which taken together indicate higher efficiency in velocity transmission from front to rear, compared to RF.

Our results indicate that fiber alignment promotes faster migration, greater persistence, and longer transmission of motion within the monolayer while randomizing fiber orientation limits all the above physical parameters. All these differences are further supported by the unjammed/jammed phenotype on AF/RF [43] as well as the differing migration front stability in both conditions. Since migratory fronts are contractile in nature and responsible for force generations in active systems, we wanted to investigate whether the observed differences in migration arise from differences in force generation and their propagations within the monolayer [16].

### Aligned fibers reduce and polarize cellular traction stresses while increasing stress cooperativity

To quantify cellular force generation, we used traction force microscopy [4]. We found that cells on AF exerted only 1/5^th^ of the total strain energy that they applied on RF, as shown in Fig. 2m. This is also evident from the traction kymograph comparison between AF and RF in x-direction (𝑇_*x*_) (Fig. 2g,h) and y-direction (𝑇_*x*_) (Fig. 2i,j) both of which reflect significantly lesser levels of forces on AF. Further, comparing 𝑇_*x*_and 𝑇_*y*_for both conditions, cells on AF apply negligible forces in the y-direction (1/5^th^) compared to the x-direction, while cells on RF apply slightly lesser forces in y-direction (0.6*𝑇_*x*_) (Fig. 2p). Thus, cell collectives on AF apply polarized forces along the direction of collagen alignment which are much lesser in magnitude compared to the directionally unbiased high pulling forces on RF. This further corroborates with our migration track analysis where cells displayed higher persistent motion along collagen alignment in x-direction. This implies, cells pulling unidirectionally along collagen fibers in x-direction need to exert minimum forces orthogonally.

**Figure 2.**
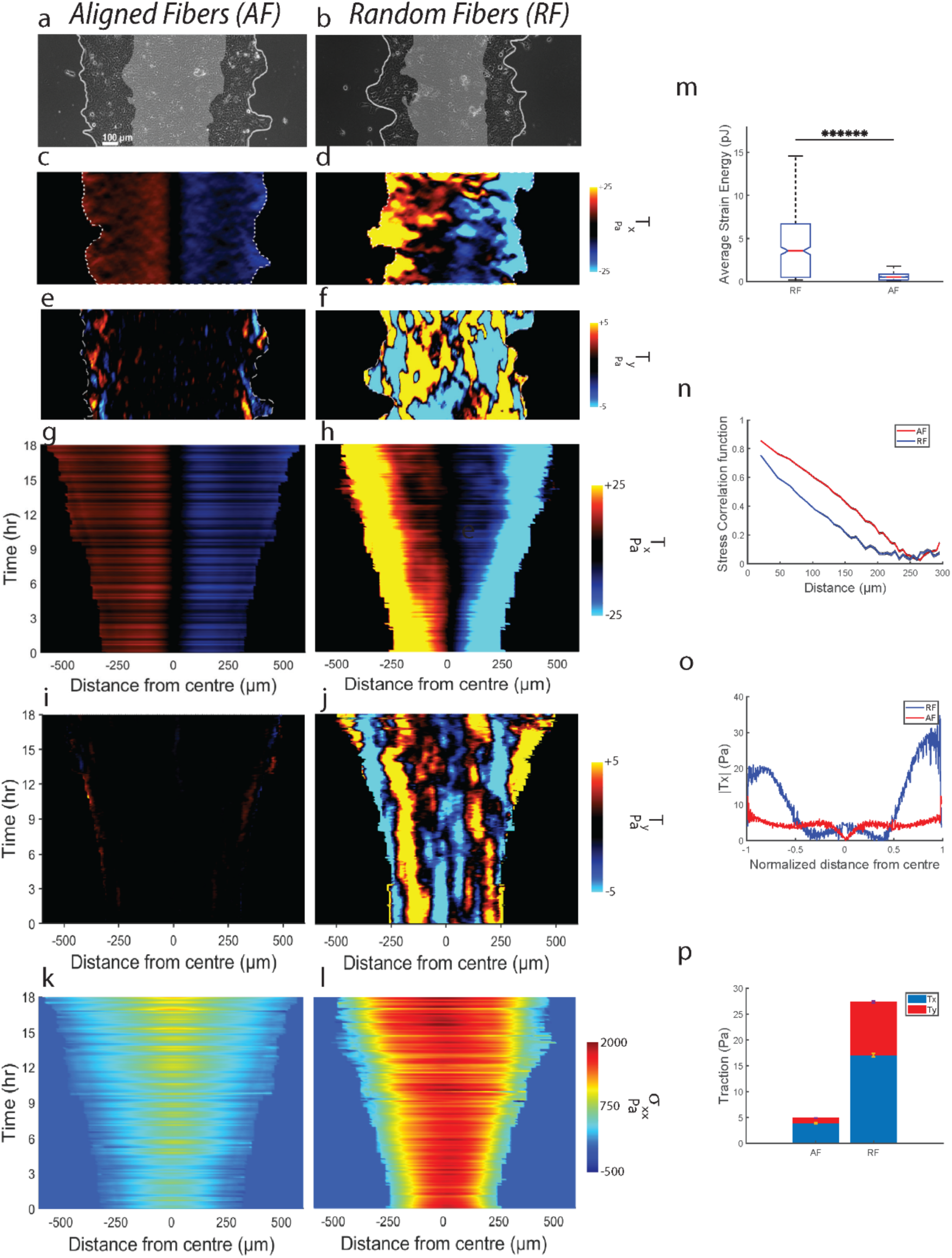
Aligned fibers cause force-effective collective migration. Representative phase contrast images of monolayer expanding on (**a**) AF and (**b**) RF. Representative heatmaps of traction component *T_x_* for (**c**) AF and (**d**) RF. Representative heatmaps of traction component 𝑇_*x*_ for (**e**) AF and (**f**) RF. Kymographs for *T_x_* on (**g**) AF and (**h**) RF. Kymographs for 𝑇_*y*_ on (**i**) AF and (**j**) RF. Kymographs for monolayer stress component 𝜎_*xx*_ on (**k**) AF and (**l**) RF. (**m**) Plot comparing strain energy between AF and RF (m=157) across n=3 (*****P=1.98×10^-84^). (**n**) Plot comparing time averaged (m=157) spatial correlation function of average-normal stresses between AF and RF (n=3). (**o**) Plot comparing time-averaged (m=157) spatial evolution of *T_x_*along the normalized width of the monolayer (n=3) between AF and RF. (**p**) Plot comparing *T_x_* with 𝑇_*y*_ for AF and RF (both *T_x_* and 𝑇_*y*_ averaged across m=157 and n=3).

We also compared force propagation across the two conditions, by measuring spatial traction distribution. While tractions were evenly distributed across the monolayer width on AF with forces equivalent to the leading-edge tractions getting exerted at 3/4^th^ distance from edge (1 being the center), there were heterogeneities in RF, with higher tractions localized only to the leading edges (1/4^th^ distance from edge), followed by a steep decrease in middle regions (Fig. 2o). This observation suggests that cell-collagen adhesions that exist deeper within the monolayer are getting strongly engaged, indicating a more effective relay of protrusive force from the edge to the center on AF compared to RF. Since relay of force occurs via myosin network attached to actin-bundles [44], [45], our analysis shows that monolayer on AF utilizes a supracellular network of actin-myosin architecture to relay forces deep within the monolayer. Further, using monolayer stress microscopy [46], we probed these differences by measuring inter-cellular stresses. From the 𝜎_*xx*_ kymographs (Fig. 2k,l), we see significantly lesser build-up of stress at the monolayer midline for AF (0.5*RF) (Fig. S2f) parallel to the traction results. To measure stress cooperativity within the monolayer, we calculated the spatial correlation function 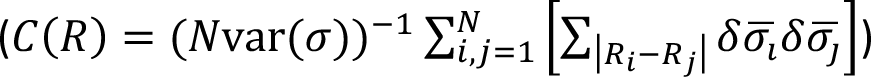 of average normal stress 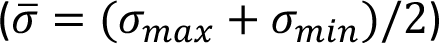. Cells on AF exhibited slower spatial decay of *C(R)*, indicating higher correlation of average normal stresses over greater distances for AF (correlation length higher by 21%). Thus, normal stresses are getting propagated deeper within the monolayer on AF. Although these AF stresses are smaller compared to RF, they are still able to cause faster expansion of monolayer both at the edge and within the monolayer.

These results are in contradiction to existing literature studying directional migration in confinement [19], aligned topographies [47], which suggest higher forces are needed for faster migration in narrower confinement or aligned nanogroove topographies. Thus, the mechanism governing collective migration on aligned fibers is distinct from those previously understood.

### Minimal shear, higher plithotaxis, and monolayer fluidization on aligned fibers explain faster migration

In conventional coupled active matter, including cell monolayers, the extent of cellular mobilization within the collective is governed by transmission of normal stresses across cell coupling junctions. Simultaneously, the local shear landscape and disruption of normal force transmission together determine local flow directions often leading to variegated collective phenotypes such as antiparallel flows [18] and vortices [19] . Since we observe greater mobilization within the monolayer along fiber alignment (AF), we next considered whether this difference between AF and RF stemmed from differences in the extent of agreement between velocities and normal stresses, a phenomenon known as ‘*plithotaxis*’ [16], and shear stresses.

To understand whether collagen alignment enhances *plithotaxis*, we rank-ordered the pairs of stress and velocity according to their distances from leading edge into quintiles and measured distribution of alignment angle *φ* between local maximal principal stress (𝜎_12/_) orientation and local migration velocity vector orientation. We find that for each quintile, the distribution of *φ* is narrower and closer to 0 degrees (Fig. 3c-f). We further calculated cumulative probability distribution of *φ* (𝑝̅(𝜑)) for each quintile. Higher alignment would be associated with a steeper rising curve for 𝑝̅(𝜑). For both AF and RF, we observe that moving away from the leading-edge causes reduction in alignment between local stress and cellular motion, indicating that *plithotaxis* is most dominant at the leading edge and progressively reduces with increasing distance from leading edge (Fig. 2r,s). However, for each quintile, AF shows stronger alignment between stress and velocity (steeper 𝑝̅(𝜑) curve), indicating more dominant *plithotaxis* compared to RF and suggests that cells are minimizing the generation of shear stress (𝜎_*xy*_) within the monolayer. We confirmed this from the 𝜎_*xy*_ kymographs (Fig. 3j,k) which show that 𝜎_*xy*_ is significantly lower on AF.

**Figure 3.**
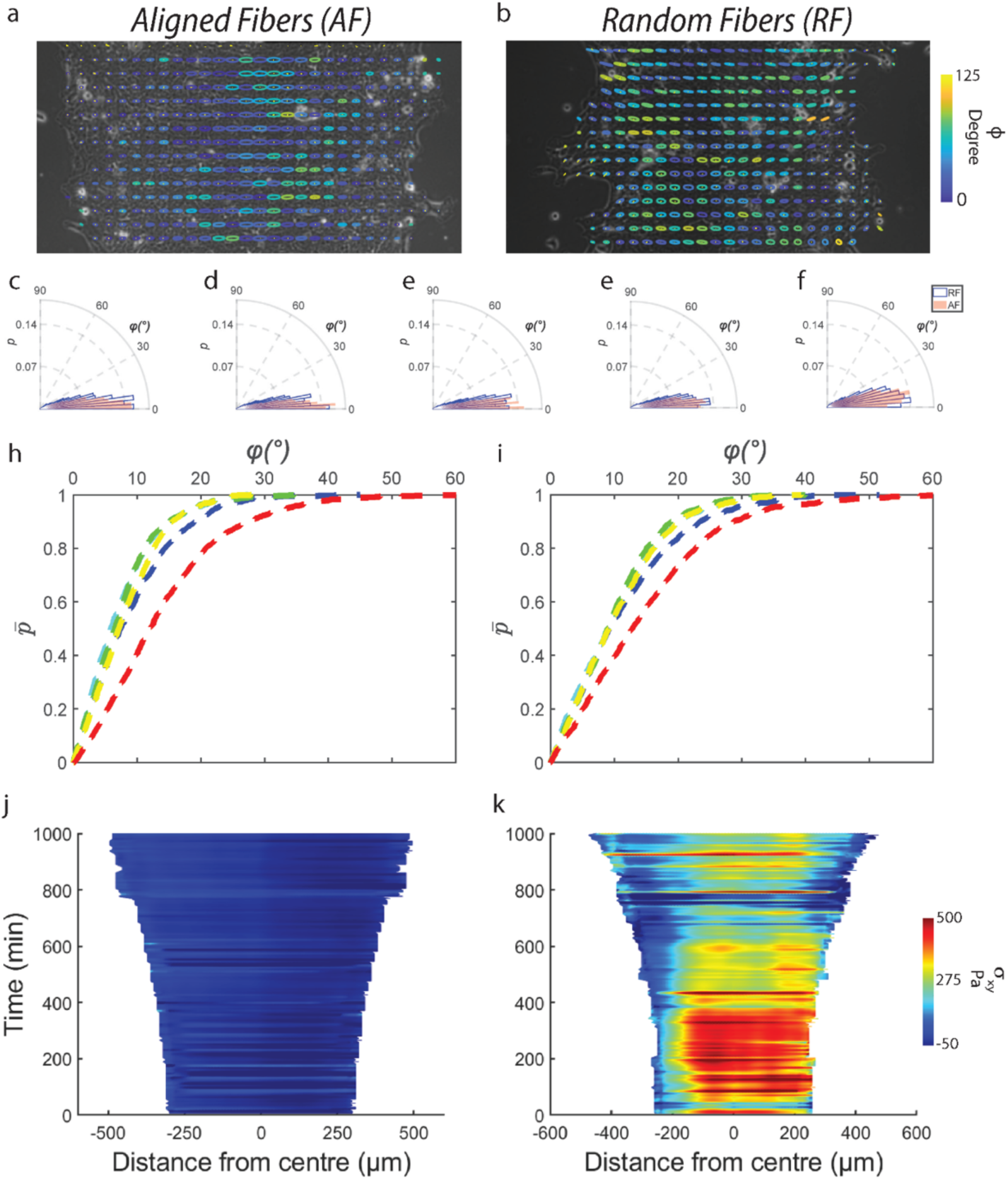
Aligned fibers cause higher *Plithotaxis* within monolayer. Representative monolayer overlaid with Principal stress ellipse which are color-coded for alignment angle 𝜑 between major axis of principle stress ellipse and direction of cellular velocity on AF (**a**) and RF (**b**). (**c**-**f**) Comparison of alignment angle 𝜑 distribution between AF and RF for quintiles based on distance from the leading edge with (c) being farthest quintile and (f) being closest quintile. Cumulative probability distribution 𝑃+(𝜑) curves (red to blue refers to quintiles at decreasing distance from leading edge, n>1000) for monolayer migrating on AF (**h**) and RF (**i**). Kymographs for monolayer shear stress (𝜎_*xy*_) for AF (**j**) and RF (**k**).

Previously, cytoskeletal fluidization has been shown to regulate dissipation of built-up stresses within cell monolayers, necessary for their active expansion [42]. Here, we measure such fluidization by measuring stress oscillations (𝜎_*xx*_) at the monolayer midline (100 µm wide strip) (Fig. 4c,d). Parallelly, we measure cellular area fluctuations (Fig. 4e,f), since recent work has shown tight coordination of cell size and stress within monolayer via extracellular signal mediated kinase activation [48]. Consistent with the strain-rate fluctuations (discussed in previous section, Fig. 4c,d), two stress peaks arise for AF within the first 10 hours, compared to a single peak in RF arising after the 10^th^ hour mark (Fig. 4c-f). Further, fluctuations in 𝜎_*xx*_ were in phase with fluctuations in cellular area (arrows in Fig. 4e-h) and at phase quadrature to strain-rate fluctuations (Fig. 4c-f)) [42]. Altogether, a higher frequency of stress dissipation on AF prevents excessive build-up of stresses within the monolayer, thus explaining why cells on AF experience reduced stresses. Frequent stress dissipation is also tied to frequent regulation of cellular areas which also explains why cell sizes on AF are smaller on AF (Fig. 1n).

**Figure 4.**
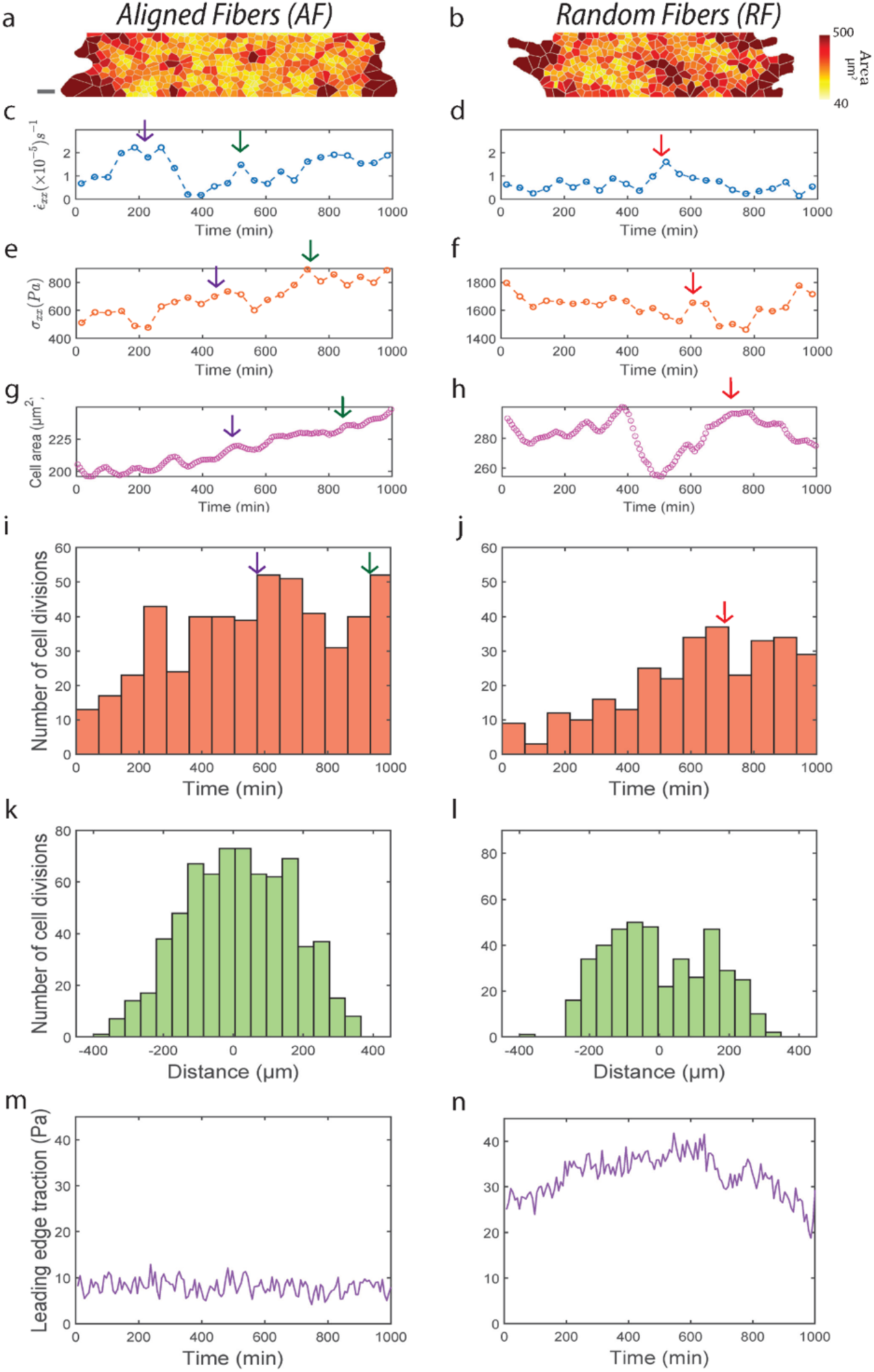
Aligned fibers cause higher fluidization within monolayer. Polygons approximating cell shapes for representative monolayer migrating on AF (**a**) and RF (**b**). Plot showing temporal evolution of 𝜀_*xx*_ (averaged across a 100 µm wide strip at the monolayer midline) on AF (**c**) and RF (**d**). Plot showing temporal evolution of 𝜎_*xx*_ (averaged across a 100 µm wide strip at the monolayer midline) on AF (**e**) and RF (**f**). Plot showing temporal evolution of cellular area (averaged across a 100 µm wide strip at the monolayer midline) on AF (**g**) and RF (**h**). Histogram showing temporal evolution of cell division frequency (averaged across a 100 µm wide strip at the monolayer midline) on AF (**i**) and RF (**j**). Histogram showing spatial distribution of cell division frequency (averaged over the entire duration of migration) on AF (**k**) and RF (**l**). Plot showing temporal evolution of leading-edge traction (averaged across a 3/4^th^ of the monolayer width) on AF (**g**) and averaged across 1/4^th^ of the monolayer width RF (**h**).

Given that stresses and cellular areas are more frequently regulated in AF, we wanted to investigate the mechanism behind this regulation. Recent literature [49]–[52] have shown that cellular stretching during monolayer expansion triggers rapid division by expediting cellular re-entry into M-phase. Thus, we measured the amount of cell divisions that occurred within the monolayer. Looking at the temporal distribution of cell divisions across AF and RF (Fig. 4 i,j), we found that cell divisions on AF were much higher than on RF throughout the 18 hours of migration and that these cell divisions were in phase with the temporal cellular area changes (arrows in Fig. 4g-j). From the spatial distribution of cell division between AF and RF, we found that cell division are maximum at the monolayer midline on AF while on RF, it is roughly 100 µm away from the monolayer center (Fig. 4k,l). The increase in cellular divisions on AF further corroborates our cell density results in (Fig. S2d) and provides an explanation for the observed differences.

From the above results, we see that AF enables strain rates within monolayer to travel more frequently and deeper within the monolayer on AF, thus transferring intra-cellular stresses deeper leading to frequent stretch-based cell division causing fluidization of the monolayer. As a result, stresses are not allowed to build-up within the monolayer allowing it to migrate at higher velocities. On RF, this chain of strain-rate and stress transfer is not as effective which would then lead to an increase in stress build-up because of infrequent fluidization causing the monolayer to stall [53].

To confirm the above ‘force effective yet fast migration’ phenotype on AF, we repeated the experiments by performing a double negative i.e., further lowering mod-PA substrate stiffness (0.6 kPa) as well as lowering the duration of fiber alignment to 4 hours (from 16 hours). Cells still migrate faster on AF, compared to RF (1.34 times) while applying much lesser strain energies (1/4^th^) on AF compared to RF (Fig. 5, S3a,c). Both cellular velocities and stresses are correlated across greater distances in clusters migrating on AF showing greater transmission of motion owing to greater force transmission (Fig. S3b,d). Expectedly, we also observed higher spatial extent of *plithotaxis* in clusters migrating on AF (Fig. S3e-i).

**Figure 5.**
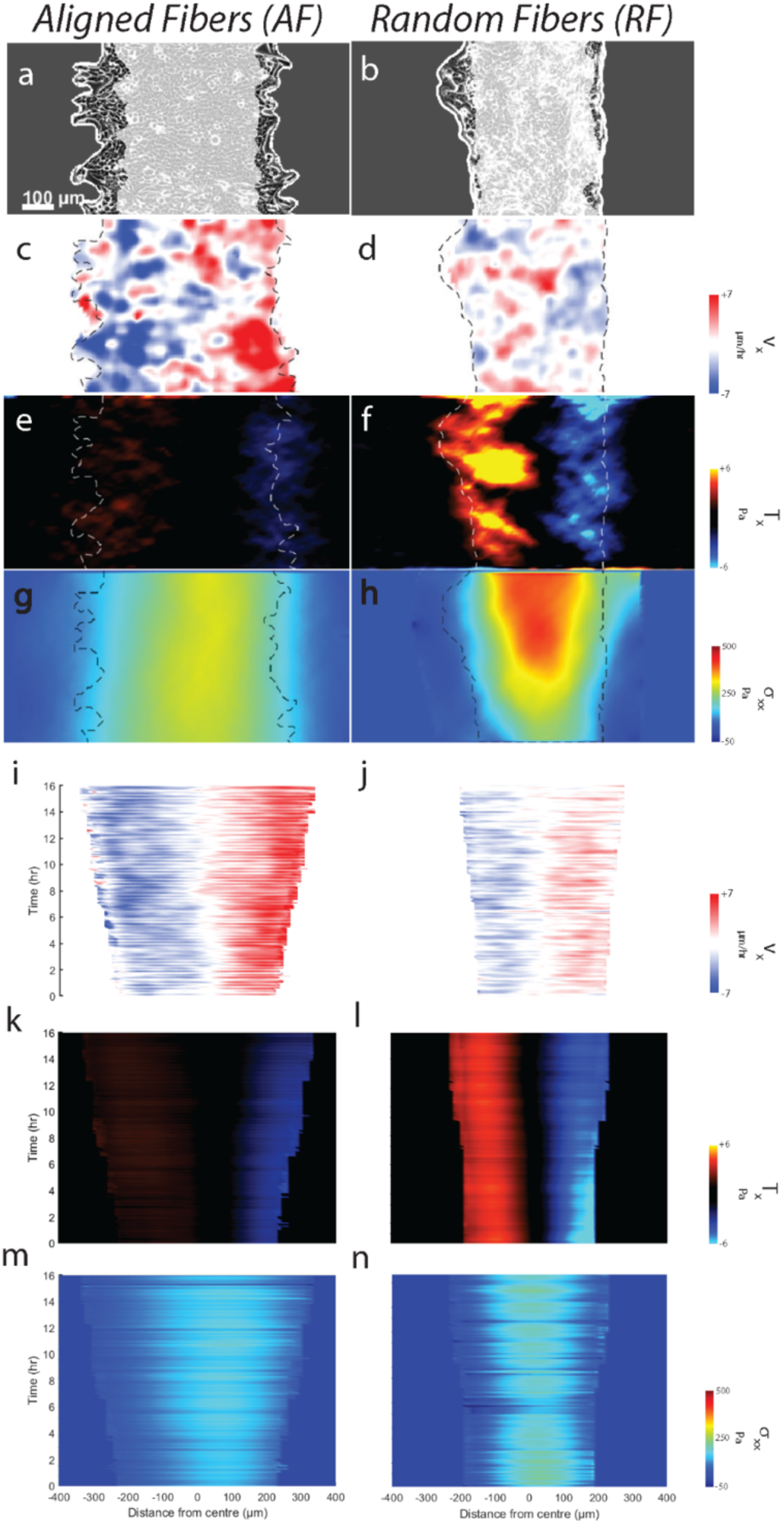
Faster force-effective migration despite reducing substrate stiffness and duration of collagen alignment. Phase contrast images of cell monolayer migrating of softer (**a**) AF and (**b**) RF. (**c**, **d**) Maps of velocity component 𝑣_*x*_. (**e**, **f**) Maps of traction component *T_x_*. (**g**, **h**) Maps of monolayer stress component 𝜎_*xx*_. Kymographs of (**i**, **j**) velocity component 𝑣_*x*_, (**k**, **l**) traction component *T_x_*, and (**m**, **n**) monolayer stress component 𝜎_*xx*_.

### Collective motor-clutch model explains ‘force-effective’ collective cell migration via rapid stabilization of cell contractility

Our experimental measurements establish that monolayers use aligned collagen fibers to expand faster while generating lesser contractile tractions and monolayer stresses. Also, we show that these forces are transmitted across greater distances within the cluster, leading to higher fluidization and subsequent release of stresses. To understand the governing mechanism behind this force-effective fast migration on AF, we developed a novel motor-clutch model in which the cell monolayer is treated as a series of Kelvin-Voigt viscoelastic springs [42] which is connected to a ligand mesh via clutch-like cell-ECM bonds [54]. The ligand mesh is connected to the substrate mesh via ligand-substrate springs mimicking covalent bonds seen experimentally [32] (Fig. 4a). In our previous paper [32], we have shown that ligands on longer collagen-1 fibers when pulled by an AFM tip, rupture higher number collagen-substrate covalent bonds compared to shorter fibers. This implies that ligands within the fibers are connected to each other, and longer collagen fibers make greater number of connections to the substrate. Since AF are both longer and aligned compared to RF (Fig. S1b-e), thus ligands on AF make greater connections to the substrate. Hence, RF is modelled as individual ligand springs disconnected from one another, while AF is modelled by connecting these ligand springs to each other, depicting ligand connectivity along the direction of migration (Fig. 4b,c). However, for both AF and RF, each substrate node offers a ligand node for binding, thus incorporating similar ligand continuity for both conditions.

We show that ligand connectivity increases migration efficiency due to greater engagement of ligand-substrate bonds when cell mesh contracts after a protrusion event, leading to shorter duration for establishing equilibrium and lesser deformation of substrate mesh (Fig. S4a-f and ‘*Single contraction experiment*’ section of supplementary text) as well as the higher rate of migration-front stabilization (Fig. S4g-j). We also show that this increased ligand connectivity mediated increase in migration efficiency is well conserved across our simulation parameters of cellular stiffness (𝑘_*c*_) (Fig. S5 a), cellular damping constant (𝜂_*c*_) (Fig. S5b) and substrate stiffness (𝑘_4_) (Fig. S5c). We also varied the extent of ligand connectivity and studied its effect on migration efficiency (Fig. S5d) and found that increasing ligand connectivity increases migration efficiency (see details in ‘*Parameter scan*’ section of supplementary text). In our model, we however see that the monolayer starts to stall after ∼5 hours of migration (Fig. S6a-b). Parallel to our experimental findings of cell division based monolayer fluidization improving migration efficiency (Fig. 4), we also implemented cell division-based fluidization in our model (see supplementary section ‘*Incorporating fluidization and spatially variation in cell stiffness and damping’* for details) and observe that monolayer does not get stalled any more (Fig. S6d-f), but the leading-edge has a convex shape indicating that with progression of time, protrusive forces produce diminished displacements at the leading-edge because the combined contractile forces due to expanded cell springs within the mesh start dominating the protrusive forces at the leading edge. This anomaly to experimental observations is addressed by inclusion of spatial variations in cellular stiffness (𝑘_*c*_), damping (𝜂_*c*_) and cell division threshold strain (𝜀_.56_) (Fig. S6j-l and Table S2) in the model (Fig. S6g-I and plot Fig. S6m).

We next compare the simulations which include fluidization and spatially varying 𝑘_*c*_, 𝜂_*c*_ and 𝜀_.56_ between connected and disconnected ligand condition (Fig. S7). We see that cells on connected ligands migrate faster (*1.7 times) (Fig. S7a-c) while applying lesser (*1.8 times) traction forces (Fig. S7d-f) than disconnected ligand condition. However, in experiments we had observed that cells on AF travel 1.56-1.33 times faster (Fig. S2a, Fig. S3a), while applying 2-3 times lower forces than RF (Fig. 2o). Furthermore, the strain-rate waves reach center first on disconnected condition (Fig. S7j,k), which is inconsistent with experiments where the X-waves reach midline first on AF (Fig. 4 c-d). Additionally, we also observe that although χ_4_ peak (Fig. S7k) is predicting unjammed motion for connected condition, it is not congruent with experimental χ_4_ peaks of AF and RF (Fig. 1 l). To address these shortcomings in our model, next we decided to include differences in actin-myosin network distribution across AF and RF which we inferred from traction distribution differences between AF and RF (Fig. 2o)), by incorporating different spatial variations in 𝑘_*c*_ (𝑥_*c*_), 𝜂_*c*_ (𝑥_*c*_) and 𝜀_.56_ (𝑥_*c*_) for connected and disconnected conditions (Fig. S8a-c)) (see supplementary section ‘*Implementing different spatial variations in stiffness, damping and fluidization across connected and disconnected ligand’* for details).

As a result, our model correctly predicts an increase in velocity (*1.38 times) in parallel with a decrease (*2.42 times) in traction forces for cells migrating on connected ligands compared to disconnected condition (Fig. 6k). Our model also correctly predicts an increase in monolayer fluidization (white arrows in Fig. 6m,o) with increase in ligand connectivity along with faster and deeper transfer of strain rate waves within the cell mesh (Fig. 6m,o). As a result, unjamming of motion emerges, as shown by the shift in χ_4_ peak to shorter time scales and reduction in peak height (Fig. 6 n), which are remarkably consistent with experiments (Fig. 1i) when compared to the disconnected condition.

**Figure 6.**
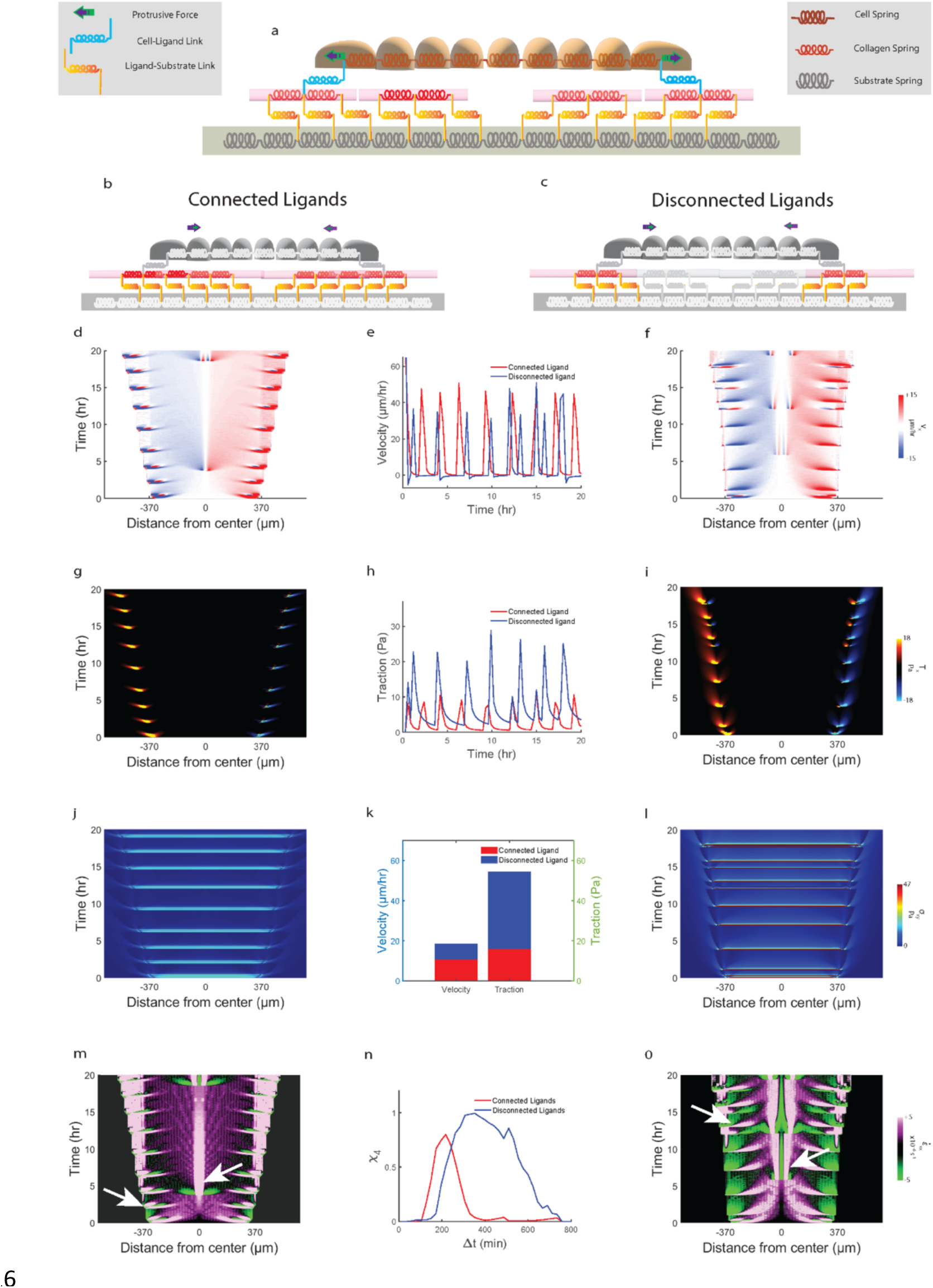
Physical model showing force-effective fast migration due to rapid contractility stabilization by fiber connectivity. (**a**) Schematic of one-dimensional model of epithelial cells represented by springs sliding viscously over rigid ligand springs fixed to substrate springs, after self-propelling forces applied at leader nodes. Higher number of ligand-substrate bonds get engaged during contractility driven retraction of monolayer in (**b**) connected ligand condition compared to (**c**) disconnected condition. Simulated kymographs of (**d**, **f**) velocity, (**g**, **i**) traction, (**j**, **l**) shear-stress and (**m**, **o**) strain-rate kymograph for connected (left) and disconnected (right) ligands. Plot comparing temporal evolution of leading-edge velocity (**e**) and traction (**h**) between connected and disconnected condition. Plot comparing average velocity and traction between connected and disconnected condition (**k**). Plot comparing four-point susceptibility (𝜒_4_) versus Δ𝑡 for simulated results (**n**).

Thus far, we have studied the effects of changing ligand connectivity on migration efficiency while keeping ligand continuity constant across the conditions. Next, we ask whether changing ligand continuity with constant connectivity has similar effects on migration efficiency. To this end, we implemented the phenomenon of *haptotaxis* using our model (Fig. 7). We implemented a gradient in ligand binding probability which restricts binding ability of cell mesh nodes to ligand nodes (equation-13, supplementary information), thus mimicking a gradient in ligand concentration along direction ofmigration (see supplementary section ‘*Implementing ‘Haptotaxis*’’ for implementation details). For all the different ligand binding gradients that we tested (Fig. 7k), we observe that steeper gradients lead to higher shift in center of mass indicating higher *haptotaxis* (Fig. 7l). However, higher gradients in ligand binding probability do not necessarily equate to higher migration efficiency, with the highest migration efficiency occurring at an intermediate ligand binding gradient (Fig. 7m). Additionally, for all the gradients tested, migration efficiency is still lower than that on connected ligand condition, indicating that ligand connectivity is more potent in terms of affecting migration efficiency compared to ligand continuity.

**Figure 7.**
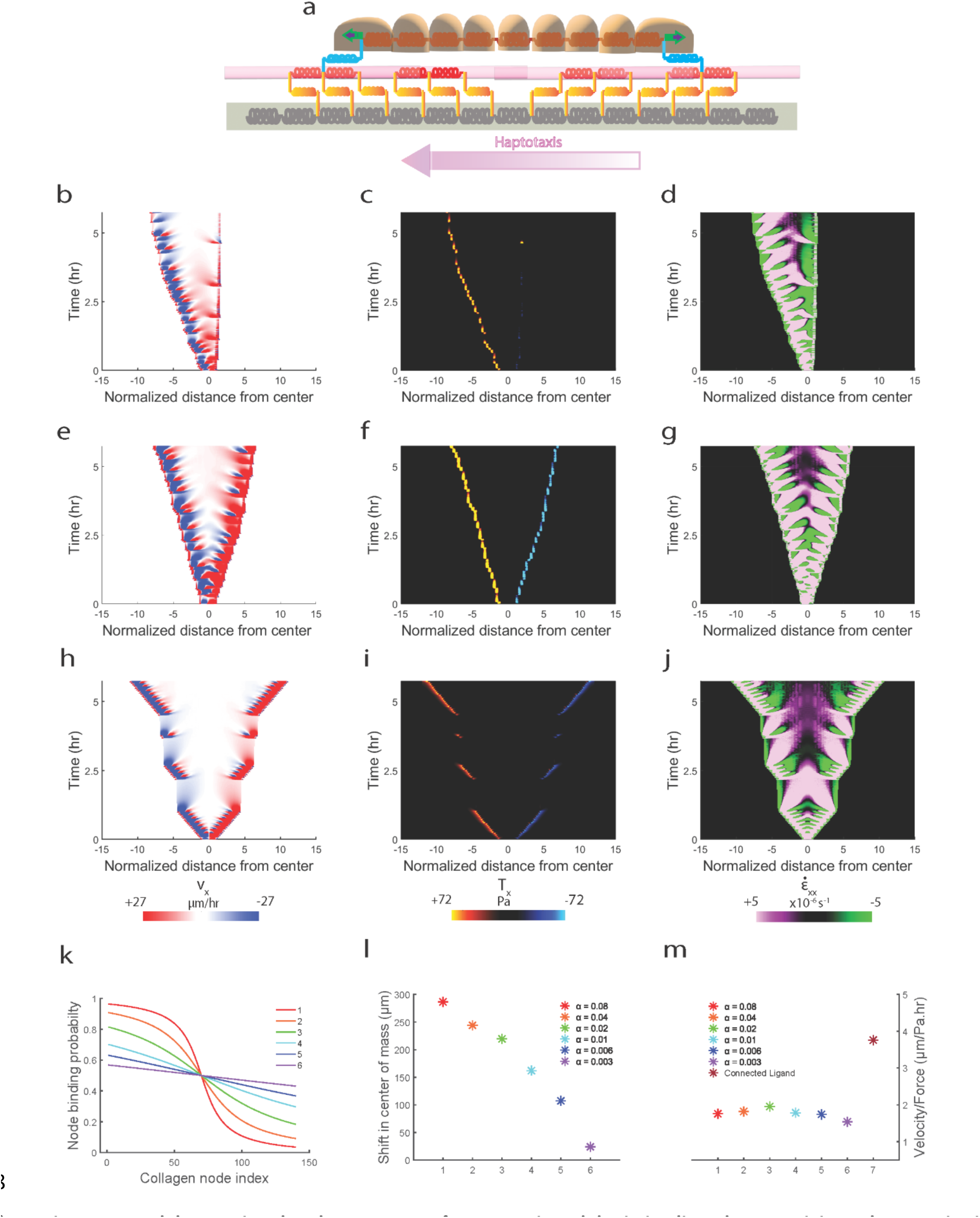
Model capturing the phenomenon of *Haptotaxis* and depicting ligand connectivity to be superior in terms of migration efficiency when compared to ligand continuity. (**a**) Schematic of one-dimensional model showing direction of increasing ligand binding probability (𝜌_*l*_) from right-to-left, discussed further in supplementary section. Simulated kymographs for (**b**) velocity, (**c**) traction and (**d**) strain-rate for 𝛼 = 0.04. Simulated kymographs for (**e**) velocity, (**f**) traction and (**g**) strain-rate for 𝛼 = 0.003. Simulated kymographs for (**h**) velocity, (**i**) traction and (**j**) strain-rate for connected ligand condition. Plot showing spatial variation of ligand binding probability (𝜌_*l*_(𝑥_*c*_)) for different values of 𝛼 (**k**). Plot showing shift in center of mass of the cell mesh migrating on ligand mesh having different 𝜌_*l*_(𝑥_*c*_) corresponding to different values of 𝛼 (**l**). Plot comparing migration efficiency for cell mesh migrating on ligand mesh having different values of 𝛼 (shown in plot k) (**m**).

Next, we sought to test whether our model could be further generalized to predict other known forms of directional migration. To this end, we decided to implement *durotaxis* – a form of directional migration towards increasing substrate stiffness, phenomenologically using our motor-clutch model. We implemented a linear gradient in substrate spring stiffness (𝑘_*s*_) (equation-14, supplementary information) with 𝑘_*s*_ increasing from left-to-right (Fig. 8a) (see supplementary section ‘*Implementing ‘Durotaxis*’’ for implementation details). We observe that implementing gradient in stiffness causes cell mesh to undergo a shift in center of mass towards increasing stiffness (Fig. 8b-d), while cell mesh migrating on uniformly stiff (Fig. 8e-g) and soft substrates (Fig. 8h-j) expand uniformly without shift in center of mass although monolayer expansion on uniformly stiff substrates is much higher than uniformly soft substrates [55]. We next tested the extent of *durotaxis* on varying stiffness gradients (Fig. 8k) and found the stiffer gradients in stiffnesses displayed higher *durotaxis* (higher shift in center of mass) (Fig. 8l) [29].

**Figure 8.**
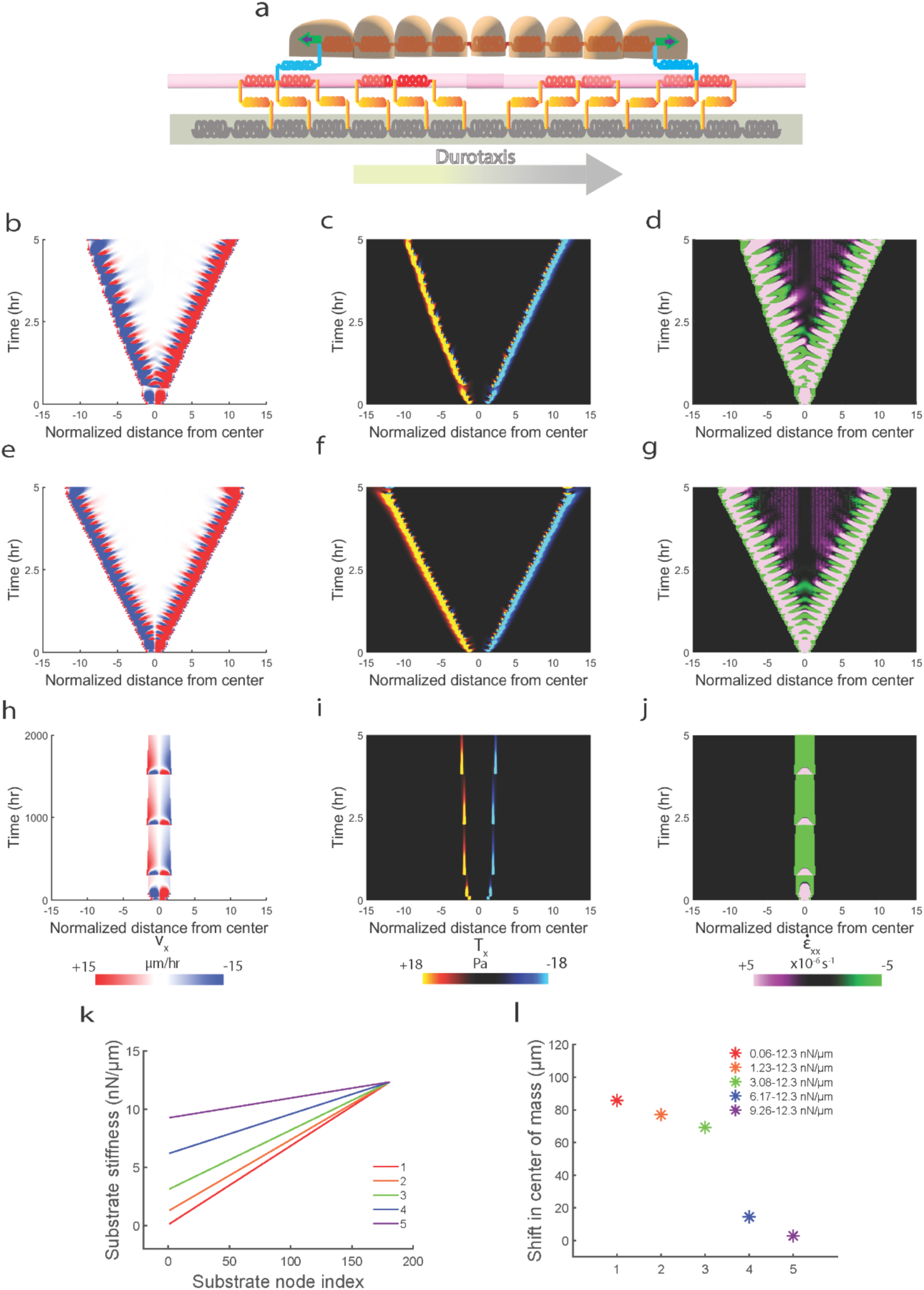
Model capturing the phenomenon of *Durotaxis* and showing higher rate of contractility stabilization is sufficient to capture the phenomenon. (**a**) Schematic of one-dimensional model showing direction of increasing substrate stiffness (𝑘_*s*_) from left-to-right, discussed further in supplementary section. Simulated kymographs for (**b**) velocity, (**c**) traction and (**d**) strain-rate for *durotaxis* stiffness range of 0.06 𝑛𝑁/𝜇𝑚 < 𝑘_*s*_ < 12.3 𝑛𝑁/𝜇𝑚. Simulated kymographs for (**e**) velocity, (**f**) traction and (**g**) strain-rate for uniformly stiff substrates (𝑘_*s*_ = 12.3 𝑛𝑁/𝜇𝑚). Simulated kymographs for (**h**) velocity, (**i**) traction and (**j**) strain-rate for uniformly soft substrates (𝑘_*s*_ = 0.06 𝑛𝑁/𝜇𝑚). Plot showing different spatial variation of substrate stiffness (𝑘_*s*_(𝑥_*s*_)) used for testing the *durotaxis* model (**k**). Plot showing shift in center of mass of the cell mesh migrating on substrate mesh having different variations in 𝑘_*s*_(𝑥_*s*_) (shown in plot k) (**l**).

## Conclusions and Outlook

Given close agreement between experiments and model predictions, we conclude that increased frictional resistance via parallel engagement of connected ligands enables a new, efficient mode of migration wherein lower net forces generate faster cell migration on matrices that offer higher ligand connectivity. While cell-cell force propagation has been shown to regulate collective migration [29], our experiments use constant matrix stiffness across conditions, thus controlling for cellular mechanotransduction, and elucidate how matrix fibers cause supracellular directional migration through long-range intercellular force transmission. We also speculate that utilizing aligned ECM fibers on 2D soft substrates gives rise to a quasi-2D mode of contact-guided directional migration. In this mode, cells utilize the connectivity of ligands offered by aligned fibers, thus delineating the role of ligand connectivity from ligand continuity presented by uniformly coated ligands on micropatterned protein stripes or topographical grooves [18], [19].

As evident from our experiments, uniform ligand coating of random collagen fibers leads to ligand continuity but not connectivity, which restricts the frictional resistance offered by ligand substrate bonds which do not get parallelly engaged as seen in case of aligned fibers. Similar phenomenon has been observed in single cells when they experience changes in ligand continuity [56], where single cells migrate faster yet apply lower forces when they experience ligand continuity compared to sharp decrease in ligand continuity. At the scale of a single cell, short fibers offer higher ligand connectivity compared to abrupt break in ligand continuity. This rise in migration efficiency mediated by aligned fiber has direct implications in cancer metastasis where tumor microenvironment presents collagen fibers of varying alignment concomitant with cancer progression. For example, recent work has shown that cell collectives use significantly less ATPs while migrating on aligned fibers compared to randomly oriented fibers [57], while breast tumor histology revealing aligned fibers offers poor prognosis [58], implying that tumor cells align matrix fibers to metastasize in an energy efficient manner. Our work reveals a previously unexplored dimension of ligand connectivity and its connection to collective migration efficiency. These findings present a new paradigm of migration efficiency regulated not just by force but by the rate of migration front stability. While only recently studies have shown the importance of migration front stability in dictating directional migration [30], [31], we show that a higher rate of front stability leads to more efficient migration and further show that this phenomenon is enough to capture *durotaxis*, a known form of efficient directional migration.

## Acknowledgements

The authors acknowledge technical assistance and equipment sharing by Dr. James Quirk for using MRI strength magnet and Professor Vijay Ramani for the use of vacuum filtration and rota-evaporator. This work was supported by the NIH/NIGMS grant R35GM128764 (to AP) and National Science Foundation, Science and Technology Centers, Center for Engineering MechanoBiology grant CMMI:154857 (to AP and MF). AP and AB conceived the project and designed study. AB performed experiments and computational modeling, analyzed data, interpreted findings, made figures, and wrote the manuscript. BS, JZ and MF contributed methods and reagents for polyacrylamide gel synthesis. AP edited the manuscript, supervised the project, and acquired funding.

