## Supplementary materials for "Fast yet force-effective mode of supracellular collective cell migration on aligned fibers due to extracellular force transmission"

### Supplementary Information

#### Materials and Methods

**Cell culture**. MCF10A cells (Tissue Culture Support Center, Washington University, St Louis, USA) were cultured in DMEM/F12 (Invitrogen) supplemented with 5% (v/v) horse serum (Invitrogen), 20 ng/ml epidermal growth factor (EGF, Miltenyi Biotec Inc), 0.5 mg/ml hydrocortisone (Sigma-Aldrich), 100 ng/ml cholera toxin (Sigma-Aldrich), 10 µl/ml insulin (Sigma-Aldrich) and 1% (v/v) penicillin-streptomycin (Sigma-Aldrich).

**Microfabrication of magnetic PDMS stencils.** Cell monolayer was confined to a particular geometry by depositing magnetic PDMS stencil with a rectangular opening (500 μm) over the gel (Fig. S1)[1]. Briefly, a mixture of PDMS and magnetite (25% w/w) was added to a mold which was 3D printed (Proto Labs Inc.). The mold containing PDMS mixture was kept in an oven to bake at 80$^{o}$ C. A magnet was attached to the bottom of the well plate just below the PA gels so that it could secure the magnetic PDMS stencil on top of the wet PA gel.

**Fabrication of mod–PA hydrogels**. Polyacrylamide (PA) gels were chemically modified by a method previously described in [2]. Briefly, N-hydroxyethyl acrylamide (HEA) was oxidized to its primary aldehyde group N-ethanal acrylamide (EA) using pyridinium chlorochromate (PCC). The crude oxidized product was first purified via vacuum filtration to remove larger impurities and further purified by column chromatography using silica as solid phase and Ethyl Acetate as the mobile phase. Briefly, crude product was dissolved in Chloroform before running it through the column to separate product which was obtained as a white flakey solid upon evaporating excess solvent in a rota-evaporator (Buchi). Fourier-transform infrared spectroscopy (FTIR) was performed to confirm the presence of aldehyde group in the purified product (Fig. S1f).

The synthesized EA was then incorporated into PA to facilitate protein conjugation. Modified Polyacrylamide gels of desired stiffness were prepared on glass-bottom 6 well-plates (Cellvis). Precursor PA solution was prepared by mixing acrylamide (A, Bio-Rad), EA and bis-acrylamide (B, Bio-Rad) (2.8% A:2.8% EA: 0.44% B) to result in 2.4 kPa stiffness gels as reported previously [2]. Softer gels were synthesized using recipe (4.592% A:1.0% EA, 0.44% B) to result in approximate stiffness of 0.3 kPa. Polymerization was initiated by adding 0.5% ammonium persulfate (APS, Sigma-Aldrich) and 0.05% tetramethylethylenediamine (TEMED, Sigma-Aldrich) to the precursor mixture. Then, 60 µL of the precursor solution is sandwiched between a hydrophobic cover slip treated with Sigmacote (Millipore) and the silanized glass bottom 6 well plates. Next, the solution was allowed to polymerize in a vacuum chamber for 45 mins.

**Magnetic alignment of Collagen-1.** The modified PA gels were then incubated in 0.1 mg ml$^{-1}$ type 1 collagen (rat tail collagen, Santa Cruz Biotechnology) inside the bore of a horizontal bore electromagnet for 2 hours at 4 C while fibillogenesis took place. Field strength for the magnet was 11 T (Fig. S1). Collagen fibrils aligned at an angle of 90$^{o}$ to the direction of magnetic field due to presence of a positive diamagnetic anisotropy of the growing collagen fibrils [3][4]. The aligned fibrils deposited onto the gel and formed covalent linkages with the modified PA gel as described previously [2].

For gels with uniform collagen deposition, modified PA gels were incubated with 0.1 mg ml$^{-1}$ type 1 collagen at 37 C for 1 h.

**Fluorescent labelling of collagen and imaging.** Collagen was fluorescently labelled according to method described in [5]. Briefly, 1.5 ml of a 3 mg/ml collagen 1 solution (pH ~7.5) was gelled at 37°C inside a 12 multiwell plate and incubated with 0.2 M sodium bicarbonate bRFfer (pH 9.0, Sigma-Aldrich) for 10 min. Next, gel was incubated in 500 µl of Sulfo-Cyanine5 NHS ester dye (Lumiprobe Co. USA) solution (in DMSO) at room temperature in dark for 1 h. Remaining NHS ester was quenched by adding 3 ml of 50 mM Tris-HCL bRFfer (pH 7.5, Sigma-Aldrich). Later, stained collagen gel was washed with PBS 6 times over a period of 2 h. Afterwards, labelled gel was solubilized using 200 mM HCl (Sigma-Aldrich). The solubilized collagen solution was dialyzed against 20 mM acetic acid (Sigma-Aldrich) at a 1:1000 ratio with continuous stirring at 4°C for 4 h. Working collagen solution was prepared by replacing 4% of the unlabeled collagen solution with its labelled counterpart. Collagen coated hydrogels were imaged using Zeiss LSM 880 laser confocal microscope (Carl Zeiss Microscopy, Germany) (Fig. S1 b-c).

**Cell monolayer patterning on soft hydrogels.** Prior to using PDMS stencils for patterning, they were stored in 70 % ethanol overnight. To prevent cell and ligand adhesion to PDMS, stencils were then passivated by incubating them in 2 % Pluronic F-127 (Sigma-Aldrich) dissolved in PBS for 1 h. During the same time, gels which were coated with collagen are washed twice in PBS and incubated with media at 37° C. Magnets were attached to the bottom of the glass bottom dishes just below the hydrogels. Passivated PDMS stencils were washed twice in PBS and air dried for 20 min, and then were deposited on the surface of the hydrogels. Next, 100 µl of media containing 60,000 cells was added into the exposed region of the hydrogel and kept inside incubator for the cells to attach. After 1 hour, unattached cells were washed away and 100 µl of fresh media was added. Twelve hours after seeding, cells reached desired cell density and stencils were carefully removed using tweezers and 3 ml of fresh media was added onto the wells.

**Time-lapse microscopy.** Migrating cells were imaged using Zeiss AxioObserver Z1 microscope (Carl Zeiss Microscopy) fitted with incubation. Cells were imaged after a 7 min interval over a period of 20 h using a 10x objective. Cells were allowed to acclimatize to the microscope incubation for 3 h before beginning image acquisition. Experiments were conducted in 37° C and 5% $CO_2$.To include the entire width of the monolayer, two images with 10% overlap were captured and then stitched using Zeiss stitching tool.

**Cell velocities and trajectories.** Velocity fields for migrating cells were computed via particle image velocimetry (PIV) though PIVlab package in MATLAB [6]. Source code was modified to include a two-dimensional Hanning window across the interrogation window to improve PIV resolution and prevent boundary artifacts during cross-correlation. Velocity field $(v_{ij})$ was obtained by running PIV for three passes of 64-, 32- and 16- pixel windows. Monolayer boundaries were computed using home-built MATLAB script. Cell trajectories were obtained by tracking labelled cell nuclei using Fiji plugin TrackMate. Cell trajectories were imported in MATLAB and further analyzed using custom written scripts. Average cellular persistence in motion was measured as $\bar{p} = \langle \frac{L_{i,e}}{L_{i,c}} \rangle$, where $L_{i,e}$ is Euclidean distance covered by the cell $i$ and $L_{i,c}$ is the contour length of its track. $\langle \ldots \rangle$ denotes average over all such tracks.

**Quantifying dynamics of cellular motion.** To quantify dynamics of cellular motion, we calculated the self-overlap order parameter [7].

$$Q(\Delta t) = N^{-1}\sum_{i=1}^{N} w_i$$

Where $N$ is the number of cells, $w$ = 1 if $|r_i(t+\Delta t) - r_i(t)| < 0.9 * d_c$ (where $r_i(t)$ is x-position of cell at time $t$ and $d_c$ is average cell diameter) and $w$ = 0 otherwise. To measure the size and lifetime of the cooperative fluctuations within the migrating cluster of cells [8], the spatially heterogeneous dynamics in cellular motion was quantified by the four-point susceptibility $\chi_4$, given by

$$\chi_4(\Delta t) = N[\langle Q(\Delta t)^2\rangle - \langle Q(\Delta t)\rangle^2]$$

Where $N$ is the number of cells, $\langle \ldots \rangle$ shows the average over sequence of images at all times, $t$. $\chi_4(\Delta t)$, is the four-point susceptibility and quantitatively shows the size of cooperatively motile cell clusters and their associated rearrangement times.

**Fourier-transform traction microscopy.** Traction forces were calculated from gel deformations which were recorded by measuring displacements in fluorescent marker beads (0.2 µm, Invitrogen) embedded inside the hydrogels on which cells were migrating. Displacements of the beads were measured in reference to the initial undisturbed condition obtained after trypsinization (10x) of cells. Bead strain field was calculated using PIV. This strain was then used to calculate the traction field by solving the inverse problem using a non-regularized Fourier transform algorithm previously employed in [9]. The algorithm was implemented in MATLAB. The total energy $U$ contribution from the monolayer towards the elastic distortion of the substrate was calculated by [10]

$$U = (\frac{1}{2})\int \vec{T}(\vec{r}).\vec{u}(\vec{r})dxdy$$

**Monolayer stress microscopy.** Stresses within the monolayer were calculated using monolayer stress microscopy implemented in MATLAB [11]. $\sigma_{ij} = \left(K_1 - \frac{2}{3K_2}\right)\dot{\varepsilon}_{kk}\delta_{ij} + 2K_2\dot{\varepsilon}_{ij}, K_1$and $K_2$ are bulk and shear viscosities, $\dot{\varepsilon}_{ij}$ is strain-rate tensor. Finite element scheme was used to model cell monolayer, assumed to behave as thin sheet of uniform thickness (Poisson's ration 0.5, 5 µm height). Boundary conditions at the leading edge were assumed to be stress free. At the top and bottom optical edges, normal displacements and shear stresses were assumed to be zero ($u_i n_i = 0, \sigma_{ij}n_j t_i = 0$) [12].

**Spatial autocorrelation function of tension.** Extent of spatial cell-cell coordination in terms of velocities and stresses was quantified by the spatial autocorrelation function [13]

$$C(R) = \frac{1}{N\mathrm{var}(\bar{\sigma})}\sum_{i,j=1}^{N}\left[\sum_{|R_i - R_j| = R} \delta\bar{\sigma}_i \delta\,\bar{\sigma}_i\right]$$

$\delta\bar{\sigma}_i$ is local departure of tension at position $R_i$ from the spatial mean $\langle\sigma\rangle$, while $\mathrm{var}(\bar{\sigma})$ is the variance of those departures, and $|R_i - R_j| = R$ denotes the bin width which consists of $N$ points. We used the same function to quantify the spatial coordination of velocity vectors. Plots were generated by averaging 4 consecutive values for smoothening purposes.

**Kymographs.** Every pixel's position within the monolayer was computed from the nearest leading edge. Next, we calculated mean values of velocities, velocity strain rates, tractions and monolayer stresses of all pixels at a given distance from leading edge. These values were then represented as a single dimension segment of the monolayer for a single time point. These segments were obtained for every time point and staked together to result in the kymographs.

**Quantifying cell shapes.** 900 cells for AF and 600 cells for RF were manually tracked over 185 frames by annotating their nuclei using MATLAB image labelling app. Labelled cell nuclei were then used to generate a Voronoi tessellation to approximate cellular boundaries. These boundaries were used to approximate the area of cells. The cell's aspect ratio (AR) was then computed from the ratio of second area moments of $I$ for the polygon [14]. Codes were implemented in MATLAB.

$$I = \begin{bmatrix} \int x^2 dA & \int xydA \\ \int xydA & \int y^2 dA \end{bmatrix}, \; AR \equiv \sqrt{\frac{I_1}{I_2}}$$

**Statistical analysis.** Statistical testing is done using a two-sample t-test. In figures, m represents the total number of time steps that were analyzed, and n represents total number of observations. Wherever n is not mentioned, it indicates one representative video. The statistical test was done using MATLAB.

#### One-dimensional motor-clutch model of monolayer expansion

***Overview.*** As shown in (Fig. 4a), the model is composed of cell monolayer, substrate and an intermediate layer of ligand which connects monolayer to substrate. Each of these layers is modelled as a collection of springs and dashpots (Kelvin-Voigt) connected in series. Connector springs between cell monolayer and ligand springs depict adhesion complexes represented by various proteins like Integrin, Paxillin, Vinculin and Talin which bind actin to extra-cellular matrix proteins [15] represented here by the ligand layer. The ligand layer is attached to the underlying substrate via links which represent the covalent attachment between collagen fibers and mod-PA gels in experiments. Additionally, the protrusive part of the monolayer is represented by protrusive nodes at each end of the cell layer which is acted upon by a constant force $F_p$. Force is transmitted across cell-cell junctions and to the substrate via intermediate ligand layer from the monolayer edges [16]. Next, we summarize the different components of the model and its numerical implementation.

***Cell monolayer.*** Monolayer is modelled as a collection of springs connected in series. While protrusions are implemented only at the leading edge, cellular contractility is represented by the contraction force which gets exerted when cell springs get stretched after protrusion. This contraction force is transferred to the substrate mesh via the ligand mesh and represents the traction force exerted by the cell monolayer mesh.

***Ligand layer.*** We implemented aligned fibers in the 1-d by joining multiple ligand nodes in series to form connected ligands. Random fibers were modelled as disconnected ligands with every two nodes disconnected from the third node leading to discontinuity in ligand connectivity (Fig. 4 b, c). Every ligand node has equal binding probability to cell node which is set to 1. We vary this binding probability spatially as discussed in '*Haptotaxis*' section later. Every ligand node is connected to substrate node via springs representing covalent linkages between collagen fiber and mod-PA gel.

***Substrate layer.***

Substrate is modelled as a series of springs connecting substrate nodes and have uniform stiffness. We vary the stiffness of substrate springs spatially while implementing '*Durotaxis*' and discuss in later section.

***Model implementation.*** Force balance at every cell node $x_{c_i}$ reads (1), where $F_{c-c}$ represents force at cell-cell junction (2) and $F_{c-l}$ represents force at cell-ligand junction (3). $F_p$ represents protrusive force and $F_{c-l}$ only act on first node and last node ($i = 1\ OR\ i = N$). Here, $\varepsilon_{c_i}$ denotes strain experienced by spring connecting nodes $x_{c_i}$ and $x_{c_{i+1}}$, and $\dot{x_{c_i}}$ denotes velocity of node $x_{c_i}$.

$$F_p + F_{c-c} + F_{c-l} + k_c\varepsilon_{c_i} - k_c\varepsilon_{c_{i+1}} - \eta_c\dot{x_{c_i}} = 0 \quad (1)$$

$$F_{c-c} + k_c\varepsilon_{c_{i-1}} - k_c\varepsilon_{c_i} = 0 \quad (2)$$

$$F_{c-l} + k_{c-l}\varepsilon_{c_i} - k_{c-l}\varepsilon_{l_i} = 0 \quad (3)$$

Force balance at every ligand node $x_{l_i}$ reads (4), where $F_{l-l}$ represents force at ligand-ligand spring junction (5) and $F_{l-s}$ represents force at ligand-substrate spring junction (6).

$$F_{l-l} - F_{c-l} + F_{l-s} + k_l \varepsilon_{l_i} - k_l \varepsilon_{l_{i+1}} - \eta_l \dot{x}_{l_i} = 0 \quad (4)$$

$$F_{l-l} + k_l \varepsilon_{l_{i-1}} - k_l \varepsilon_{l_i} = 0 \quad (5)$$

$$F_{l-s} + k_{l-s} \varepsilon_{c_i} - k_{l-s} \varepsilon_{l_i} = 0 \quad (6)$$

Force balance at every substrate node $x_{s_i}$reads (7), where $F_{s-s}$ represents force at substrate-substrate spring junction (8).

$$F_{s-s} - F_{l-s} + k_s \varepsilon_{s_i} - k_s \varepsilon_{s_{i+1}} - \eta_s \dot{x}_{s_i} = 0 \quad (7)$$

$$F_{s-s} + k_s \varepsilon_{s_{i-1}} - k_s \varepsilon_{s_i} = 0 \quad (8)$$

For each incremental time step, the following algorithm is implemented:

1. Apply protrusive motor force to leader node only if it is at equilibrium (at rest).
2. Solve equations 1, 2 and 3 for nodes representing the cell mesh.
3. Solve equations 4, 5 and 6 for nodes representing the ligand mesh.
4. Solve equations 7 and 8 for nodes representing the substrate mesh.
5. The displaced cell leader node attaches to the nearest ligand node engaging clutch.
6. Check strain for every cell and perform cell division if strain is greater than the threshold division strain.

***Single contraction experiment***

To see whether forces exerted by the contracting monolayer are different across connected and disconnect ligand conditions, we studied the cell mesh contraction after a single protrusion event. In (Fig. S4 c), we observe that cell mesh contracting on connected ligand comes to rest within 6 mins while for disconnected ligands, cell mesh comes to rest only at 21 mins (Fig. S4 d). Also, the cell mesh undergoes lesser displacement at the leading edge while contracting in case of connected condition. This is also shown by the traction plots (Fig. S4 e, f) where the substrate mesh on connected ligand condition undergoes lower contraction due to cell mesh getting stabilized quicker resulting in lesser traction forces compared to disconnect ligand condition. When we repeat protrusion event for connected condition, we see that cells can make three protrusions in the same time cells make one on disconnected ligand condition (Fig. S4 g, f) and for each protrusion, the resulting contractility driven tractions are lesser in magnitude to the traction for the single event taking place on disconnected ligand condition (Fig. S4 i, j). Mathematically, we can explain the above phenomenon as follows. In Fig. S4 (a and b), when we consider force balance for ligands alone,

$$F_{cell-ligand} = \sum_1^N F_{ligand-substrate} \quad (9)$$

Clearly, $N$ is greater for connected ligand condition (Fig. S4 a) when compared to disconnected ligand condition (Fig. S4 b). When we consider the fiber (red box) containing ligand (red spring), then $F_{ligand-ligand}$ becomes an internal force and will cancel out. Only external forces $F_{cell-ligand}$ and $F_{ligand-subtrate}$ should play a part in the force balance. Also springs connecting ligands are much stiffer than springs connecting ligand to substrate or cell to ligand ($k_{ligand-ligand} \gg k_{ligand-substrate}$, $k_{ligand-ligand} \gg k_{cell-substrate}$). Thus, fiber is unlikely to deform due to external force $F_{cell-ligand}$, meaning ligand springs undergo minimum compression/expansion.

Now, when cells apply $F_{cell-ligand}$ while contracting after a protrusion event, in case of connected ligand condition, this force is balanced by greater number of $F_{ligand-substrate}$. As a result, substrate gets deformed lesser, and the contracting cell node comes to rest quicker. $F_{cell-ligand}$ is same across the two conditions since leader nodes experience same protrusion forces ($F_p$).

***Parameter scan***

Next, we let the simulation for both the conditions run for greater time steps. We then performed a scan across the different parameters that exist in our model to check whether the above phenotype of faster migration and lower traction forces on connected ligands is conserved for the range of parameter values (Fig. S5). We first vary the stiffness of cell springs ($k_c$). In Fig. S5 a, we observe that for all the stiffnesses tested, cells on connected ligands travel faster yet apply lesser traction forces compared to disconnected ligand condition represented by the ratio of average leading-edge velocity to the average leading-edge traction forces. A higher ratio indicates higher migration efficiency. For each condition, we however observe that increasing stiffness of the cell springs reduces migration efficiency. Since stiffness of cell springs is a measure of cell contractility, this plot shows that increasing contractility without increasing protrusive forces can negatively affect collective migration efficiency implying that there needs to be balance between protrusive and contractile forces for effective collective migration [17]. We next varied the cellular damping constant ($\eta_c$) (Fig. S5 b), and observe that for all tested values, migration on connected ligand is more efficient than disconnected ligands, although increasing cellular damping reduces efficiency, implying that friction due to maturing cell-cell contacts leads to jammed amorphous motion [18]. Next, we varied substrate stiffness by varying the stiffnesses of individual springs ($k_s$) comprising the substrate mesh (Fig. S5 c) and we observed that for all tested values, connected ligand condition displayed higher migration efficiency compared to disconnected condition. We also observe that increasing substrate stiffness increases migration efficiency [19]. Next, to measure the effect of ligand connectivity on migration efficiency in the disconnected condition, we varied the number of ligand nodes that were connected to each other. We observe that increasing ligand connectivity increases migration efficiency (Fig. S5 d). We however observe, that across all the tested parameters, the monolayer's expansion stalls after ~5 hours. We will tackle this problem next.

***Incorporating fluidization and spatially variation in cell stiffness and damping***

Since the cell mesh begins to stall after ~5 hours of expansion, it indicates that the combined contractile forces within the mesh due to the expanded cell springs are dominating the protrusive forces at the leading edge, preventing the monolayer from further expansion. This also results in a convex curvature of the leading edge with progression of time arising from lower displacements upon protrusion because of increase contraction within (Fig. S6 a-c). From experiments we have observed that cells undergo active fluidization within the monolayer to dissipate built-up stresses which in-turn allows the monolayer to further expand (Fig. 4). Hence, we implement cell-division based fluidization within the monolayer, where cell springs divide when the strain reaches a threshold amount. We decided to incorporate a spatial variability in the ability of cells to divide following our experimental observations that cell divisions occur most frequently near the middle of monolayer rather than at the edges (Fig. 4 k, l). Thus, we implement a spatially varying threshold division strain ($\varepsilon_{c_{th}}(x_c)$) according to equation 10 (Fig. S6 j).

$$\varepsilon_{c_{th}}(x_c) = \alpha_\varepsilon + (\gamma_\varepsilon) * \left(\delta_\varepsilon - \frac{1}{\sigma_\varepsilon\sqrt{2\pi}} e^{-\frac{1}{2}\left(\frac{x_c-\mu_\varepsilon}{\sigma_\varepsilon}\right)^2}\right) \tag{10}$$

Upon implementation of fluidization in our model, we see that cells are now able to able to continue expansion beyond the 5-hour mark (Fig. 6 d-f). However, we still observe the convex shape of the leading edge with a brief period of stalling (between 8 and 13 hours) which discontinues once the forward propagating strain rate wave which originated at the center following cell division reaches the leading edge (Fig. S6 f). After which, the monolayer continues to protrude for the rest of the simulation duration (24 hours). Since the middle of the monolayer experiences maximum stretching and given that recent experiments have shown that leader cells have much higher stiffness compared to follower cells [20], we decided to implement a spatial variation in cell stiffness ($k_c(x_c)$) within the monolayer (equation 11) with higher spring stiffness near the edges and softer springs at the center following distribution in Fig. S6 k.

$$k_c(x_c) = \gamma_k(\alpha\omega_k + \delta_k\beta_k - \frac{1}{\sigma_k\sqrt{2\pi}} e^{-\frac{1}{2}\left(\frac{x_c-\mu_k}{\sigma_k}\right)^2}) \tag{11}$$

Parallelly, we implemented a spatial variation in cellular damping constant ($\eta_c(x_c)$) (equation 12), inverse with respect to cellular stiffness ($k_c(x_c)$) variation (Fig. S6 l), following the rationale that stiffer cells would have higher actin organization leading to more elastic characteristics (quicker deformations) compared to softer cells which would have delayed response to stress transfer, but more deformable hence viscous [21]. Additionally, since cell crowding due to proliferation occurs at the center, this leads to higher friction causing contact inhibition of motion compared to leading edges where there are more open spaces and lesser cellular densities. Thus, to capture 2-D contact inhibition in 1-D, the damping constant follows equation 12.

$$\eta_c(x_c) = \beta_\eta + \gamma_\eta(\frac{1}{\sigma_\eta\sqrt{2\pi}} e^{-\frac{1}{2}\left(\frac{x_c-\mu_\eta}{\sigma_\eta}\right)^2} - \delta_\eta) \tag{12}$$

As a result of these additions to the model (parameters in Table S2), we see that the cells are now able to expand without stalling and the kymograph leading edge develops a concave morphology (Fig. S6 g-i) like those seen in experiments (Fig.1 e-f). We also see that including fluidization and spatial variations in cell stiffness ($k_c(x_c)$) and damping ($\eta_c(x_c)$) further improve the migration efficiency for both connected and disconnected condition but for each condition, connected ligands still have higher efficiency than disconnected condition (Fig_S6 m).

Next, using the above spatially varying cell stiffnesses, damping and threshold strain for division (Table S2), we run simulations for both connected and disconnected conditions (Fig. S7). From the velocity kymographs (Fig. S7 a, c), we see that cells expand faster on connected fibers (also shown in leading-edge velocity comparison in plot b). Also, cells apply lesser forces on connected fibers as seen in traction kymographs (Fig. S7 d, f) and leading-edge traction comparison plot (Fig. S7 e) while developing lesser shear stresses within the cell layer (Fig. S7 g, i). However, cells on connected ligand condition travel twice as fast compared to disconnected condition, while applying tractions two times lesser on the same. We have seen in our experiments, that cells apply as much as two to three times lesser traction forces on AF (Fig. 2 o, p and Fig. 4 m, n) while travelling 1.33-1.56 times faster on the same compared to RF (Fig. S2 a, Fig. S3 a). Additionally, there are discrepancies in the strain-rate propagation between the simulated and experimental observations. We see that strain-rate propagation reaches the center on disconnected ligand (before 7-hour mark) faster than connected ligand (around 8-hour mark) (Fig. S7 j and k), which is contrary to what we observed in experiments (Fig. 1 g and h) and (Fig. 4 c and d). Additionally, we also observe that although $\chi_4$ peak (Fig. S7 k) is predicting unjammed motion for connected condition, it is not congruent with experimental $\chi_4$ peaks of AF and RF (Fig. 1 l). We next address these shortcomings in our model.

In plot (Fig. 2 o) comparing average traction exerted across the monolayer width (x-direction) between AF and RF, we see that cells on AF apply steady forces from edge to center of monolayer while on RF, high forces are only localized at the edges while monolayer interior exerts minimal forces. For cells to exert comparable tractions at the center and edge on AF indicates that focal adhesions are mature throughout the width of monolayer and capable of generating leading-edge level high tractions while on RF, mature focal adhesion exist only near the leading edges while in the interior of the monolayer only weak adhesions exist as indicated by the drastic tapering of traction forces. Since myosin proteins attached onto actin fibers cause contractile forces within monolayer [22][23], the spatial differences in traction distribution indicates differences in actin-myosin distribution between AF and RF. Thus, in addition to differences in ligand connectivity in our model, we next incorporate differences in actin-myosin contractility to further improve our model predictions.

***Implementing different spatial variations in stiffness, damping and fluidization across connected and disconnected ligand***

Parallel to differences in experimental leading-edge traction distribution on AF and RF (Fig. 4 m, n), we incorporate differences in actin-myosin contractility between connected and disconnected ligand condition in our in-silico model. Since in our model, stiffness of the cell spring ($k_c$) is a measure of actin-myosin contractility, we decided to incorporate variability between the spatial variation of ($k_c(x_c)$) in connected and disconnected condition (discussed previously in Fig. S6 g-i) by incorporating different spatial variation on cell spring stiffness ($k_c(x_c)$) for connected and disconnected condition as shown in Fig. S8 a. We implemented inverse differences in spatial distribution of cell damping constant ($\eta_c(x_c)$) (with respect to ($k_c(x_c)$), following the assumption that uniform spatial cellular stiffness on connected ligand condition would imply uniform spatial cellular viscosity and vice-versa on disconnected ligand condition (Fig. S8 b). Additionally, we also incorporate differences in spatial variation of threshold division strain between connected and disconnected ligand condition (Fig. S8 c) given our observation of differences in position of maximum cell divisions between AF and RF (Fig. 4 k, l) and previous literature showing that cell stiffness modulates cell-division frequency [24][25]. Using these differences in parameters (enumerated in Table S3), we run simulations (Fig. 6).

##### *Implementing 'Haptotaxis'*

In our model, the differences between connected ligands and disconnected ligands occurs due to the differences in amount of ligand connectivity between the two conditions. This is shown in Fig. S5 plot (d), where increasing the ligand connectivity leads to an increase in migration efficiency. However, in both our simulation conditions, the amount of ligand continuity is constant, i.e., for both the conditions, there are equal number of ligands for the leading-edge cell node to attach onto as it protrudes outward. We wanted to know if changing ligand continuity also has a similar effect on migration efficiency like ligand connectivity. *Haptotaxis* is a form of chemotaxis, where cells undergo directional migration upon exposure to gradients in ligand concentrations. In this phenomenon, emphasis is on changing the number of attachment-points that cells are exposed to as opposed to changing the interconnectivity of the attachment points, which we have studied so far. Thus, to test the effect of ligand continuity on migration efficiency, we implement *haptotaxis* using our model (Fig. 7). To this end, we introduce a ligand binding probability ($\rho_l$) in our model which limits the node binding capacity of the leading cell edge node. We thus mimic scarcity, or abundance of ligands by decreasing or increasing ligand binding probability. Spatial variation of ligand binding probability ($\rho_l(x_c)$) follows equation 13. We further varied $\alpha$ to obtain multiple variations for $\rho_l(x_c)$ (Fig. 7 k) and measured extent of *haptotaxis* for each condition (Fig. 7 l-m).

$$\rho_l(x_c) = (tan^{-}(\alpha(-x_c) + 1.5)/3) \qquad (13)$$

##### *Implementing 'Durotaxis'*

In our model, the cell layer engages in crosstalk with the substrate layer via the ligand layer, as shown in Fig._S5 plot (c), where increasing substrate stiffness ($k_s$) improves migration efficiency. We next wanted to test whether our model can simulate *durotaxis*, a phenomenon where cells directionally migrate towards increasing substrate stiffness [26][22], to show broad applicability of our in-silico model (Fig. 8). To this end, we incorporate a linear spatial gradient in substrate spring stiffness ($k_s(x_s)$) according to equation 13. Further, we varied $k_{min}$ to obtain multiple curves for $k_s(x_s)$ (Fig. 8 k) and measured extent of *durotaxis* for each condition (Fig. 8 l).

$$k_s(x_s) = k_{min} + ((k_{max} - k_{min})/x_{max})(x_s) \qquad (14)$$

##### *Numerical implementation*

Computational simulations were numerically implemented using the forward Euler method. All equations were implemented in a C++ code developed by the author. Multiple class definitions were borrowed from CHASTE library [27], an online code repository for cell-based simulations.

***Model limitations***

We have implemented focal adhesions only at the leading edges in our model. As a result, we see that in our simulation results (Fig. S7 d, f, Fig. 6 g, i), tractions are restricted to the leading edges for both connected and disconnected conditions. However, from our experiments, we clearly see that focal adhesions are getting engaged deep within the monolayer for AF while on RF, focal adhesions are restricted at the leading edges as indicated from the spatial evolution of $T_x$ (across the monolayer width – x-direction) (Fig. 2 o). We decided to implement the focal adhesions only at the edges for both ligand arrangements because of the lack of experimental measurements on the binding and unbinding rates for inner focal adhesions across AF and RF. However, we hypothesize that the differential engagements of focal adhesions across monolayer depth on AF and RF leads to differential viscoelastic properties on the cells across both the conditions. Deeper engagement of focal adhesions across the depth of monolayer on AF would lead to uniform stiffness and viscosity across the depth while on RF, higher focal adhesions at the leading edges (reflected by higher tractions at the edges and tapering tractions from thereon) would indicate high stiffness at the monolayer edges, followed by tapering elasticity and increasing viscosity as we move towards the monolayer center. We have tried to capture these aspects (consequence of differential engagement of focal adhesions) by implementing varying viscoelasticity of cells across monolayer depth on connected and disconnected condition (Fig. S8 a-c). We have roughly tried to keep the net elasticity in the monolayer for both connected and disconnected ligand condition same since cell spring elasticity in our model approximates the f-actin amount. Given that substrate stiffness is known to epigenetically regulate the nuclear architecture (by regulating relative euchromatin and heterochromatin compaction levels) thereby regulating gene expression and protein production (for example YAP) [28], [29]. Since substrate stiffness across AF and RF is same, we hypothesize that total amount of f-actin on AF and RF would be similar, however that amount gets redistributed differently dependent on the focal adhesion arrangement. Deeper focal adhesion engagement would mean longer (uniform) supracellular f-actin assembly (concentration) across the monolayer depth on AF while focal adhesion restricted to leading edges on RF would indicate shorter but more concentrated f-actin assembly at the leading edges.

***Simulation parameters.***

| ***Parameter*** | ***Symbol*** | ***Value*** |
|---|---|---|
| Protrusive force | $F_p$ | 3083 nN [30] |
| Cell spring stiffness | $k_c$ | 4.92 nN $\mu m^{-1}$ [30], [31] |
| Cell damping constant | $\eta_c$ | 0.55 nN $\mu m^{-1}$ s (Note A) |
| Ligand spring stiffness | $k_l$ | 18.51 nN $\mu m^{-1}$ |
| Ligand damping constant | $\eta_l$ | 0.22 nN $\mu m^{-1}$ s(Note A) |
| Substrate spring stiffness | $k_s$ | 6.17 nN $\mu m^{-1}$ |
| Substrate damping constant | $\eta_s$ | 0.22 nN $\mu m^{-1}$ s(Note A) |
| Cell-ligand spring stiffness | $k_{c-l}$ | 30.86 nN $\mu m^{-1}$[30] |
| Ligand-substrate spring stiffness | $k_{l-s}$ | 12.34 nN $\mu m^{-1}$ |

Table S1: Base set of parameters for simulations. For running parameter scans (Fig. S5), the specific parameter was varied (x-axis) while keeping all other parameter values constant.

| ***Symbol*** | ***Value*** |
|---|---|
| $\omega_k$ | 0.398 |
| $\delta_k$ | $1.75*10^{-71}$ |
| $\sigma_k$ | 10 |
| $\gamma_k$ | 1000 |
| $\alpha_\varepsilon$ | 0.675 |
| $\delta_\varepsilon$ | $6.11*10^{-5}$ |
| $\gamma_\varepsilon$ | 2 |
| $\sigma_\varepsilon$ | 5 |
| $\beta_\eta$ | 0.55 |
| $\delta_\eta$ | $7.89*10^{-3}$ |
| $\gamma_\eta$ | 75 |
| $\sigma_\eta$ | 10 |

Table S2: Parameters for spatial variation of $\varepsilon_{c_{th}}$, $k_c$ and $\eta_c$ (equations 10-12) resulting in plots in Fig. S6 (j, k, l) and simulations depicted in Fig. S7.

| ***Symbol*** | ***Connected ligand*** | ***Disconnected ligand*** |
|---|---|---|
| $\omega_k$ | 0.398 | 0.079 |
| $\delta_k$ | $1.75*10^{-71}$ | $1.22*10^{-4}$ |
| $\sigma_k$ | 1 | 5 |
| $\gamma_k$ | 1000 | 10000 |
| $\alpha_\varepsilon$ | 0.675 | 0.675 |
| $\delta_\varepsilon$ | $1.02*10^{-19}$ | $6.11*10^{-5}$ |
| $\gamma_\varepsilon$ | 0.8 | 2 |
| $\sigma_\varepsilon$ | 2 | 5 |
| $\beta_\eta$ | 0.55 | 0.28 |
| $\delta_\eta$ | $1.75*10^{-711}$ | $3.97*10^{-3}$ |
| $\gamma_\eta$ | 20 | 400 |
| $\sigma_\eta$ | 1 | 8 |

Table S3: Parameters for spatial variation of $\varepsilon_{c_{th}}$, $k_c$ and $\eta_c$ (equation 10-12) resulting in plots in Fig. S8 (a, b, c) and simulations depicted in Fig. 6.

**Note A:** Damping constant ($\eta_c$) used in our theoretical model is lower than the models we have referred to [30], [31]. In our model, the innovation lies in the cell node being connected to a ligand mesh which in-turn is connected to substrate mesh with all three meshes undergoing crosstalk to shape migration. Changes to ligand mesh (Fig. 6, Fig. S7, Fig. 7) can cause changes to migration of cell mesh, while changes to substrate mesh also significantly affects cell mesh migration (Fig. 8). The cell node experiences sufficient resistance from both underlying mesh apart from the contribution of the viscous resistance, thus values of $\eta_c$ are lower to accommodate resistance from the underlying meshes, an aspect not present to similar extent in the referred models. Additionally, even though value of $\eta_c$ is small, it has significant impact on migration efficiency as shown in plot (Fig. S5 b) where we test a range of values for $\eta_c$.

### Supplementary Figures:

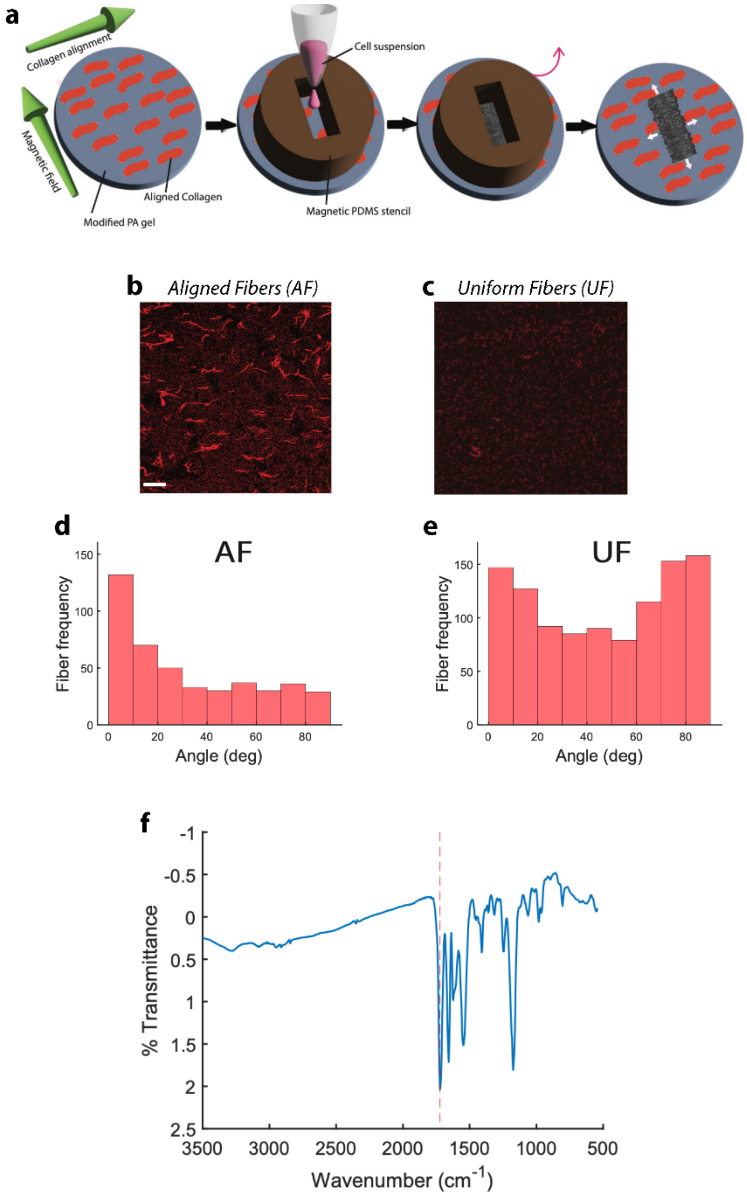


**Figure S1.** (**a**) Modified soft polyacrylamide (mod-PA) gels are exposed to high strength magnetic field to orient collagen in particular direction. A PDMS stencil is deposited on the mod-PA gel. MCF-10A cells are seeded and allowed to attach only by the gap defined by PDMS stencil. When cells reach confluency, stencil is lifted, and cells start invading available space. Representative images of fluorescently labelled collagen-1 on AF (**b**) and RF (**c**). Scale bar is 50 µm. (**d** and **e**) Histogram depicting orientation angle distribution for collagen fibrils on AF (d) and RF (e). (**f**) FTIR plot showing presence of aldehyde group peak at 1722 $cm^{-1}$. More than 400 fibers were analyzed for plots in **d** and **e**.

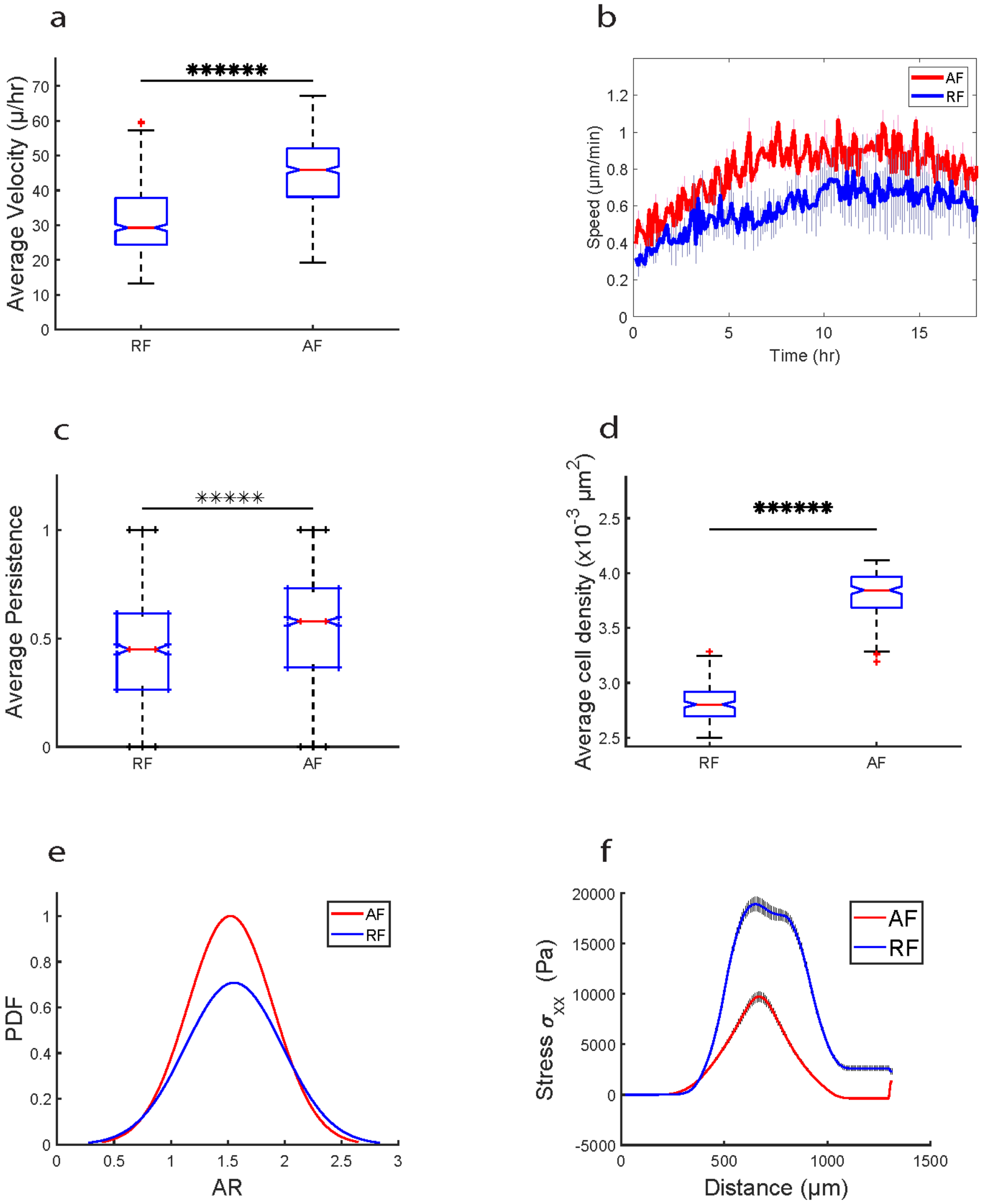


**Figure S2.** (**a**)Box-plot comparison of average velocity comparison for cells migrating on AF and RF (m = 157, n = 3, p = 1.98 x $10^{-84}$). (**b**) Average velocity of cells migrating on AF and RF as a function of time. Error is represented as standard error (m = 157, n = 3). (**c**) Time averaged cellular persistence over the entire duration of experiment (900 tracks for AF, 500 tracks for RF, p = 5 x $10^{-18}$). (**d**) Time average (m = 158) cellular density comparison for cells on AF and RF (p = 1.8 x $10^{-127}$). (**e**) Distribution of cellular aspect rations for cells on AF (red) and RF (blue). (**f**) Plot showing monolayer stress component $\sigma_{xx}$ comparison between AF (red) and RF (blue) across the width of monolayer (x-direction) and averaged over the monolayer height (y-direction) (m = 157, n = 3).

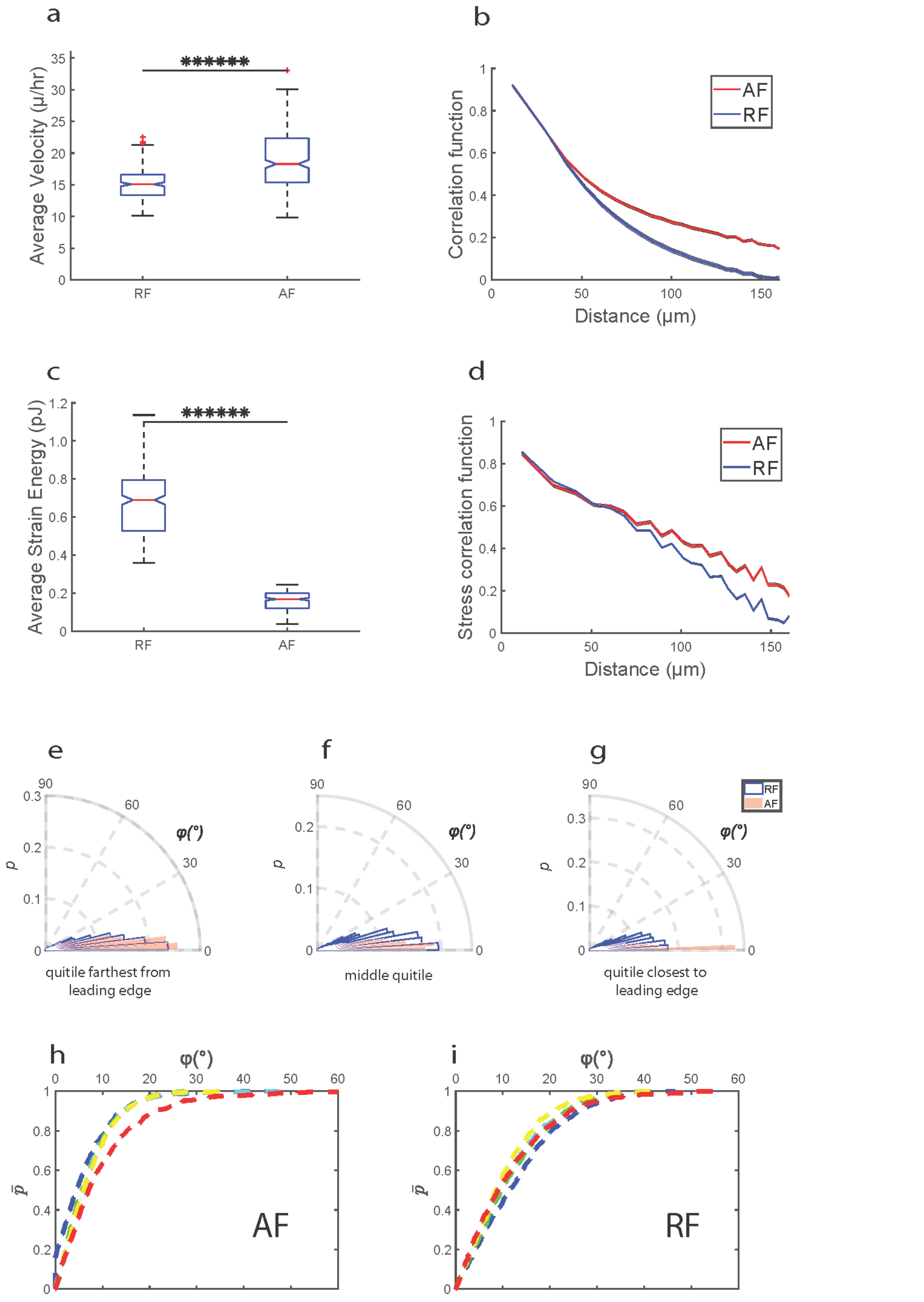


**Figure S3.** (**a**)Box-plot comparison of average velocity comparison for cells migrating on AF and RF (m = 146, n = 2, p = 1.40 x $10^{-28}$). (**b**) Time-averaged spatial autocorrelation function of $v_x$ for AF (red) and RF (blue), error is represented as standard error. (**c**) Strain energy imparted by monolayer on AF and RF averaged across entire duration of migration (m = 146, n = 2, p = 6.43 x $10^{-222}$). (**d**) Time averaged (m =

157) spatial correlation function of average-normal stresses in AF (red) and RF (blue), error is represented as standard error for n = 2 observations. (**e**-**g**) The alignment angle $\varphi$ comparison between AF and RF at quintiles furthest (e), mid-distance (f) and closest (g) to the leading edge. In all three cases, distribution is narrower for AF indicating *plithotaxis* is more dominant in AF compared to RF. (**h**) Cumulative probability distribution $\bar{P}(\varphi)$ curves, from red to blue are at decreasing distance from leading edge for monolayer migrating on AF. (**i**) Cumulative probability distribution for monolayer migrating on RF.

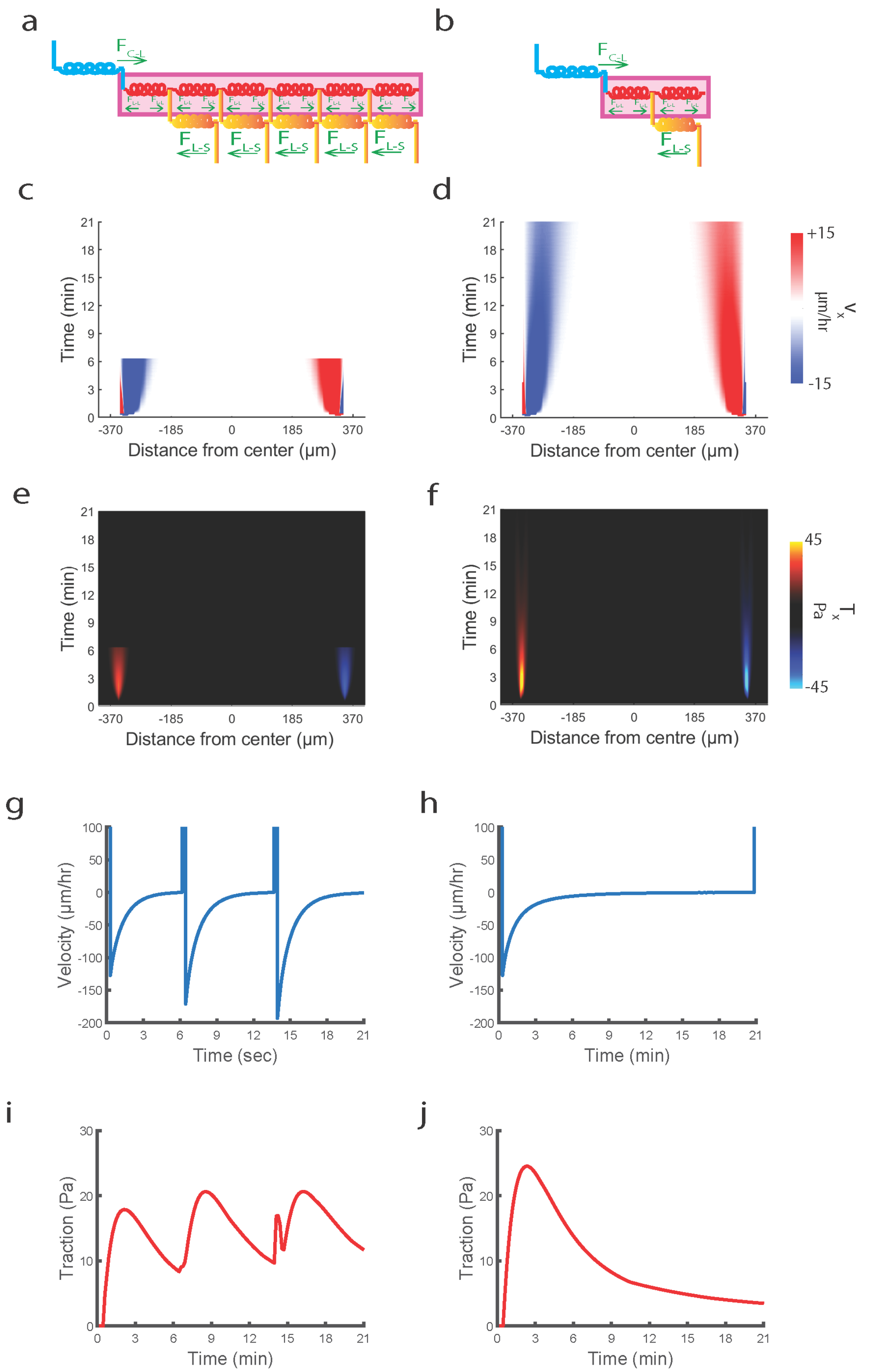


**Figure S4. Single contraction experiment.** Schematics showing differences in force balance among cell-ligand forces ($F_{c-l}$), ligand-ligand forces ($F_{l-l}$) and ligand-substrate forces ($F_{l-s}$) between connected ligand (**a**) and disconnected ligand (**b**). Forces are depicted as green arrows. Simulated kymographs of (**c,d**) velocity, (**e,f**) traction for connected (left) and disconnected (right) ligands for single contraction experiment. Plots showing velocity evolution at the leading edge for connected fiber (**g**) during the single contraction event for disconnected ligand (**h**). Plots showing traction evolution at the leading edge for connected fiber (**i**) during the single contraction event for disconnected ligand (**j**).

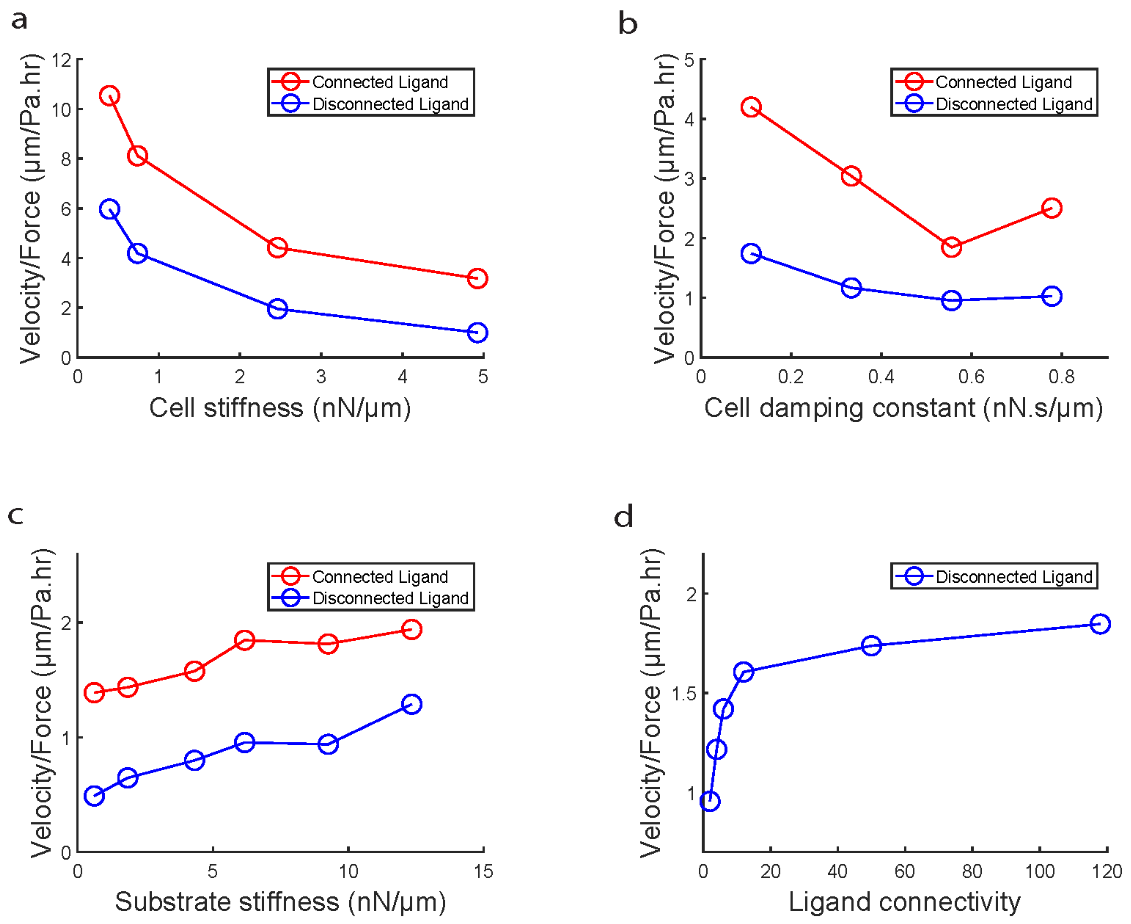


**Figure S5. Parameter scan.** (**a**) Plot comparing migration efficiency (velocity gained/traction exerted) versus cell stiffness across connected and disconnected ligand. (**b**) Plot comparing migration efficiency (velocity gained/traction exerted) versus cell damping constant across connected and disconnected ligand. (**c**) Plot comparing migration efficiency (velocity gained/traction exerted) versus substrate stiffness across connected and disconnected ligand. (**d**) Plot showing evolution of migration efficiency (velocity gained/traction exerted) versus varying ligand connectivity for disconnected ligand condition. Migration efficiency was measured for the initial 5 hours of migration for all data points.

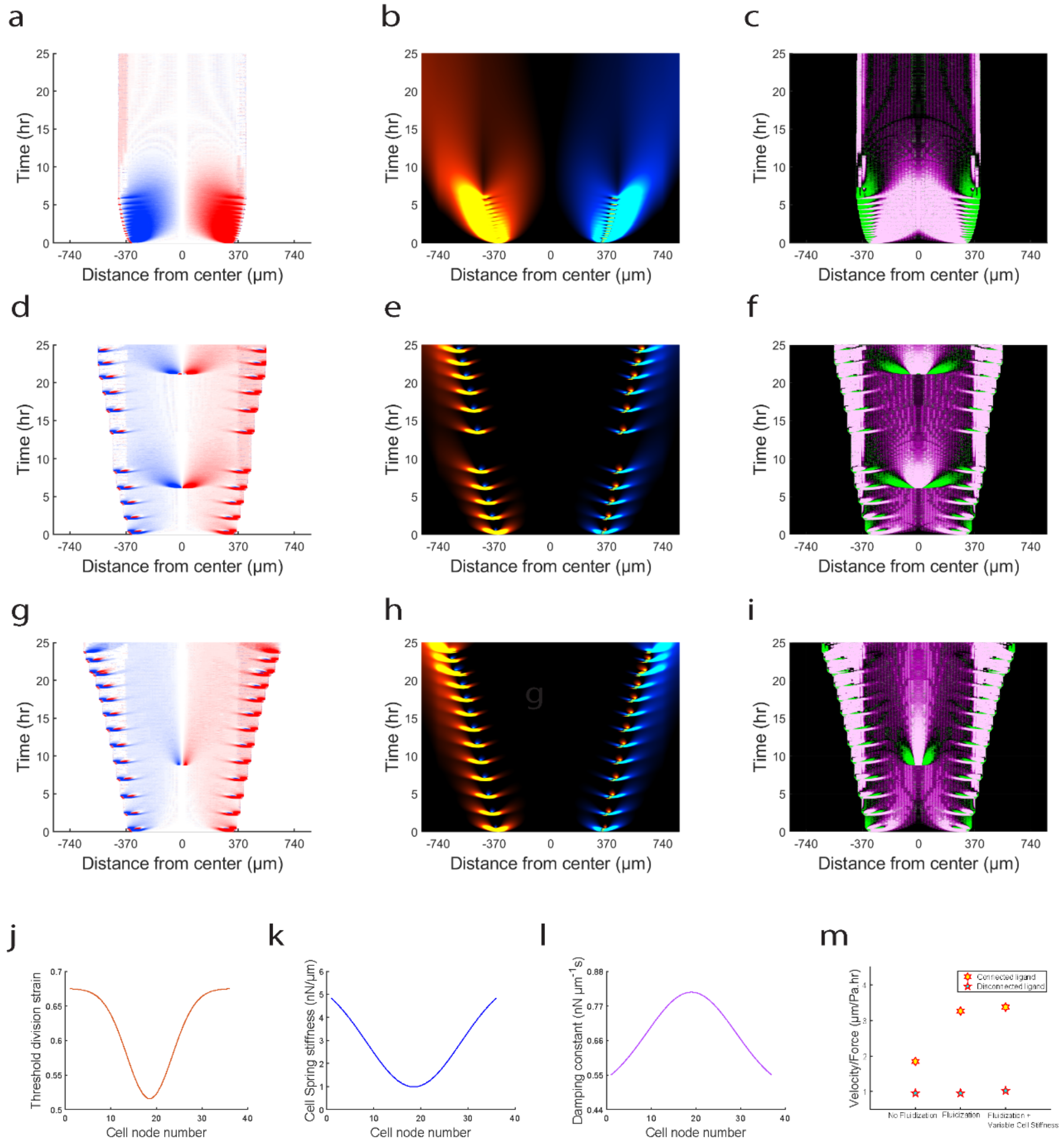


**Figure S6. Fluidization.** Simulated kymographs of (**a**) velocity, (**b**) traction and (**c**) strain-rate for connected ligand condition without fluidization. Simulated kymographs of (**d**) velocity, (**e**) traction and (**f**) strain-rate for connected ligand condition with fluidization. Simulated kymographs of (**g**) velocity, (**h**) traction and (**i**) strain-rate for connected ligand condition with fluidization and spatial variation in cell stiffness and damping constant. Plot showing spatial variation in threshold division strain across the cell nodes in silico (**j**). Plot showing spatial variation in spring stiffness across the cell nodes in silico (**k**). Plot showing spatial variation in damping constant across the cell nodes in silico (**l**). Plot comparing migration efficiency (velocity gained/traction exerted) for '*no fluidization*', '*fluidization*' and '*fluidization with spatial variation in stiffness and damping*' across connected and disconnected ligand conditions (**m**).

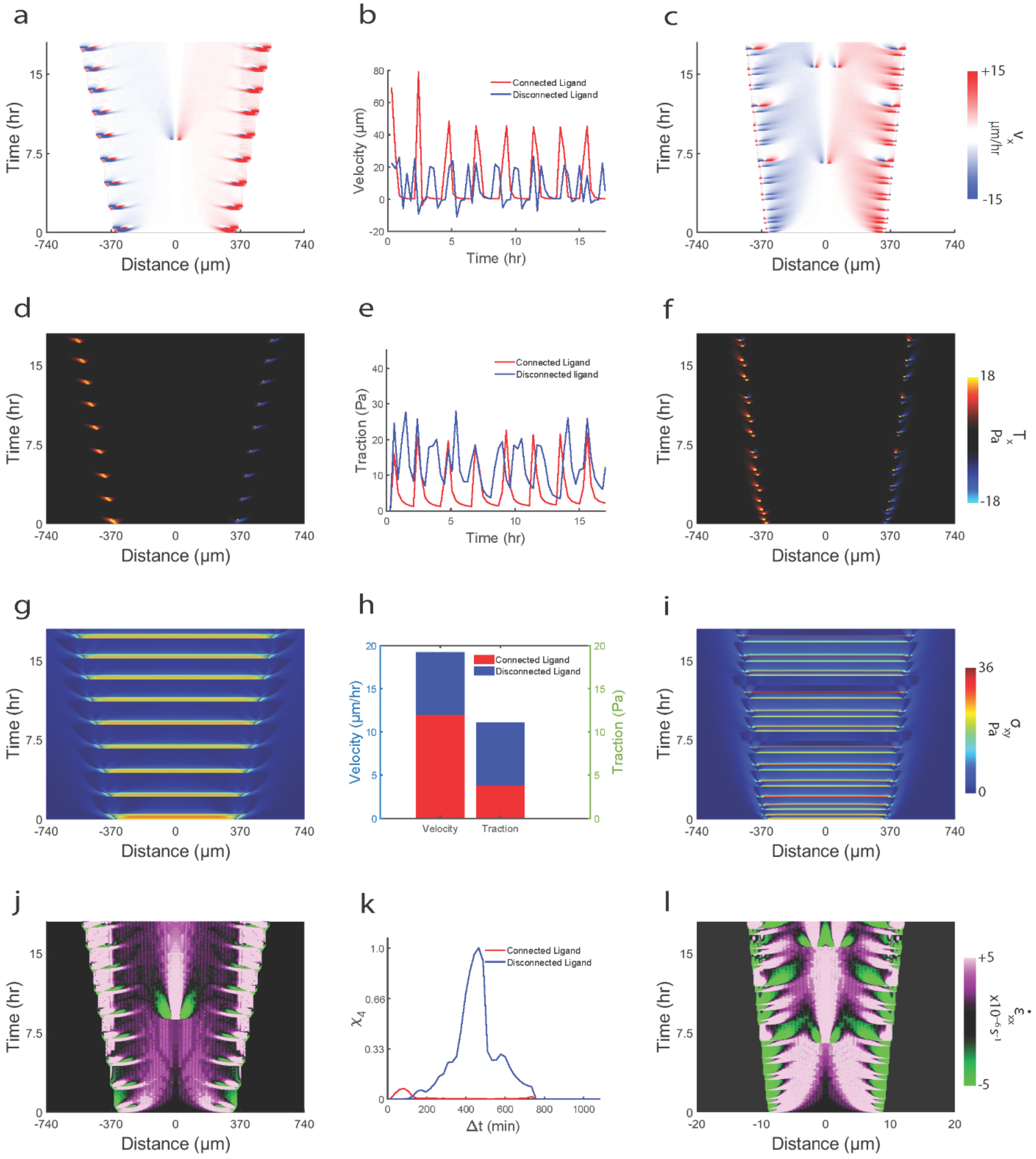


**Figure S7.** Simulated kymographs of (**a**,**c**) velocity for connected (left) and disconnected (right) conditions for '*fluidization with spatial variation in stiffness and damping'* condition. Plot comparing temporal evolution of leading-edge velocity between connected and disconnected ligands (**b**). Simulated kymographs of (**d**,**f**) traction for connected (left) and disconnected (right) conditions for '*fluidization with spatial variation in stiffness and damping'* condition. Plot comparing temporal evolution of leading-edge traction between connected and disconnected ligands (**e**). Simulated kymographs of (**g**,**i**) shear stress for connected (left) and disconnected (right) conditions for '*fluidization with spatial variation in*

*stiffness and damping'* condition. Plot comparing average leading-edge velocity between connected and disconnected ligands (left) and average leading-edge traction velocity between connected and disconnected ligands (right) (**h**). Plot comparing four-point susceptibility $\chi_4$ versus $\Delta t$ between connected and disconnected ligand conditions (**k**).

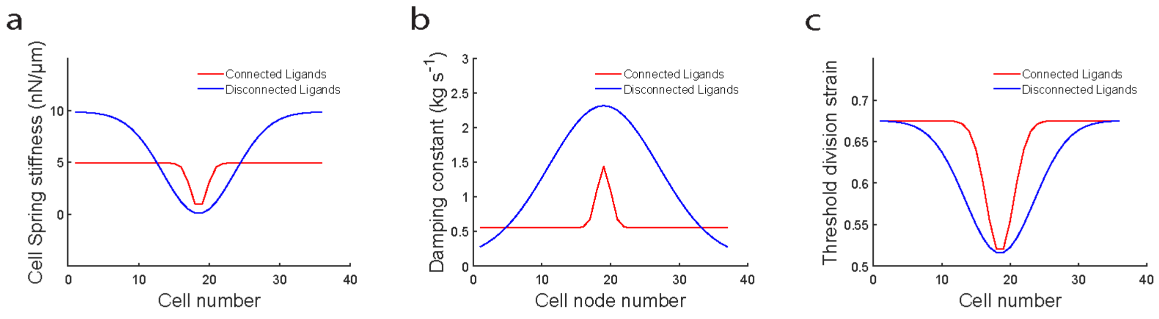


**Figure S8.** Plot comparing spatial variation in spring stiffness across the cells between connected and disconnected ligand conditions (**a**). Plot comparing spatial variation in damping constant stiffness across the cell nodes between connected and disconnected ligand conditions (**b**). Plot comparing spatial variation in threshold division strain across the cells between connected and disconnected ligand conditions (**c**).
